# Resting fMRI functional connectivity reflects fluctuations in inhibitory interneuron activity

**DOI:** 10.64898/2026.06.25.734567

**Authors:** Daniel Zaldivar, Lea Ives, Kenji W. Koyano, Rebecca Bhik-Ghanie, Brian E. Russ, Frank Q. Ye, David A. Leopold

## Abstract

Primate brain function relies on distributed cortical networks. These networks are commonly identified through fMRI functional connectivity, defined as the spatial correlation of hemodynamic fluctuations measured at rest. To assess the contribution of distinct neuronal populations to fMRI functional connectivity, we obtained concurrent fMRI and dense single-unit recordings at rest in the macaque. Then, using standard waveform-based classification of action potential shape, we compared the activity of different neural subtypes to the local and brain-wide patterns of fMRI activity. Putative excitatory neurons were functionally intermixed, with approximately half having positive and half negative correlation with the local fMRI signal. By contrast, all putative inhibitory interneurons were positively correlated, with one subclass exhibiting brain-wide correlation that closely matched conventional seed-based functional connectivity. These findings indicate that, although excitatory projection neurons may underpin long-range network communication interneuron activity most closely matches the fMRI fluctuations at the heart of resting functional connectivity.

## Introduction

The brain’s patterns of intrinsic neural activity during periods of quiescence have been used to study its interconnections and network architecture. Human functional MRI (fMRI) studies use spontaneous hemodynamic correlations at rest, often called functional connectivity, to characterize the brain’s large-scale circuit organization in health and disease (Acero-Pousa et al., 2025; Arsenault et al., 2018; Biswal et al., 1995a; Fox and Raichle, 2007; Friston, 2011; Gutierrez-Barragan et al., 2022; Huber et al., 2021; Hutchison et al., 2013b). In animal studies, spatiotemporal coupling among electrophysiological signals has similarly been used to probe functional organization across multiple spatial and temporal scales (Fukushima et al., 2012; Kodama et al., 2018; Logothetis, 2008; 2015; Stringer et al., 2019; Tsodyks et al., 1999).

The classical approach to studying functional connectivity begins with a seed, typically a fMRI voxel or cluster of voxels, whose time course is correlated with that of other voxels across the brain. The functional connectivity maps generated using this method have been used to characterize resting-state networks (Biswal et al., 1995b; Smitha et al., 2017), study disease processes (Du et al., 2018; Taylor et al., 2021; Woodward and Cascio, 2015), and elucidate large-scale network architecture (Biswal et al., 2010; Coletta et al., 2020; Gutierrez-Barragan et al., 2024). Although higher-order, data-driven methods such as independent component analysis are often preferred for capturing complex, distributed patterns of connectivity (Iraji et al., 2023; Yang et al., 2020), seed-based approaches remain important as they provide an intuitive and spatially specific measure of how activity patterns in a particular region relate to broader brain dynamics, facilitating both hypothesis-driven analyses and cross-species comparison (Coletta *et al*., 2020; Gutierrez-Barragan *et al*., 2024; Pagani et al., 2023).

The neuronal basis underlying fMRI correlations remains poorly understood, including both the anatomical and physiological factors that contribute to correlational networks (He et al., 2008; Khader et al., 2008; Leopold and Maier, 2011; Li et al., 2022; Schölvinck et al., 2013; Turchi et al., 2018; van den Brink et al., 2016; Vincent et al., 2007; Zaldivar et al., 2018). The multiple neural subpopulations that coexist in each voxel further convolute the relationship between neural underpinnings and fMRI correlations (Goulas et al., 2019; Logothetis, 2008; Young et al., 2013; Zaldivar *et al*., 2022). A subset of these neurons is presumed to carry the signals shared across areas. Through a host of potential mechanisms, their collective action on the local vascular responses ultimately translates into local fMRI responses.

A common interpretation of functional connectivity is that it reflects, at least in part, the activity of long-range excitatory projection neurons mediating interareal communication (Ding et al., 2024; Marik et al., 2010). Optogenetic stimulation of such neurons are known to elicit fMRI signal change in downstream brain areas (Lee et al., 2010). Comparison of spontaneous fMRI correlations across the brain can often show strong correspondence with anatomical connectivity assessed directly or indirectly (Honey et al., 2009; Matsui et al., 2019; Matsui et al., 2011; Vincent *et al*., 2007; Wang et al., 2013). The large-scale structure of distributed fMRI networks would seem to suggest that they reflect activity generated or synchronized through long-range anatomical connections.

On the other hand, local inhibitory neurons exert strong control over cortical excitability and vascular responses (Krawchuk et al., 2020; Tremblay et al., 2016; Uhlirova et al., 2016; Vo et al., 2023; Vo et al., 2025). Their contributions to large-scale functional connectivity are likely to be indirect, through their action on excitatory projection neurons. This hypothesized division of labor is evident in computational models of resting-state activity (Deco et al., 2011; Honey *et al*., 2009; Schmidt et al., 2018). The contribution of distinct neural subtypes to this process is an active area of investigation (Moon et al., 2021; Moon et al., 2025). Evidence from human MRI spectroscopy suggests that regional GABA concentration is inversely associated with network connectivity (Kapogiannis et al., 2013; Markicevic et al., 2020; Pijnenburg et al., 2019; Stagg et al., 2014), while local chemogenetic inhibition in animal models has led to increases in FC (Elorette et al., 2024; Rocchi et al., 2022; Sastre-Yagüe et al., 2026). Elucidating the relative contribution of neuronal cell types to the emergence and sustenance of functional network organization can vastly improve our mechanistic understanding of brain physiology in health and disease.

The present study investigates this issue by simultaneously measuring cortical single-unit activity and spontaneous fMRI fluctuations across the brain in the unanesthetized macaque. Using a standard waveform-based classification to distinguish cell types (Henze et al., 2000; Trainito et al., 2019), we compare the activity of putative excitatory and inhibitory neurons to local hemodynamic responses. Applying a seed-based correlation approach, we investigate which neural subtypes show the greatest concordance with the spatial patterns of functional connectivity across the brain. We report that narrow waveform neurons, namely putative interneurons, exhibited the strongest correlation with both local fMRI signals and brain-wide functional connectivity. All narrow waveform neurons were positively correlated with the local fMRI signal, whereas approximately half of broad waveform neurons were negative correlated. These findings indicate that fMRI measures of spontaneous activity and functional connectivity are most closely tied to the activity fluctuations of inhibitory interneuron subpopulations with a given cortical voxel.

## Results

To enable simultaneous measurement of electrophysiological and fMRI signals, we implanted MRI-compatible chronic microwire electrodes in the cerebral cortex of five macaques (Bondar et al., 2009; Koyano et al., 2021; Zaldivar et al., 2022). In each case, the microwire tips were clustered within ∼1 mm^3^ volume, approximating the size of a single fMRI voxel. The recording sites were in the inferior temporal cortex, targeted for previous studies based on their fMRI face selectivity (Koyano *et al*., 2021; Zaldivar *et al*., 2022). These regions have been previously shown to exhibit anatomical interconnections with other face-selective cortical regions (Grimaldi et al., 2016; Moeller et al., 2008).

During the experiments, the animals sat quietly in darkness in the scanner bore. In sixty-seven 30-min, we recorded single-unit and local field potential activity concurrent with fMRI scanning (**Fig. 1A** and **S1**, see Methods) (Liu et al., 2018; Schölvinck et al., 2010; Zaldivar *et al*., 2022). We evaluated the spiking activity of 157 unique isolated single units, often exploiting the stability of the microwire recordings to combine individual cells’ data across sessions and days (Koyano et al., 2023; Zaldivar *et al*., 2022). A total of 157 unique, well-isolated neurons were recorded, including 79 from the anterior fundus face patch (36 neurons in M1, 30 in M2, and 13 in M3) and 78 from the anterior medial face patch (34 neurons in M4 and 44 in M5). Time-varying fluctuations in instantaneous firing rate were computed based on spike counts measured during gap periods built into the fMRI sequence to be free of MRI artifacts (Zaldivar *et al*., 2022) (see Materials and Methods). Data from some of the recording sessions were analyzed in a previous study that focused on the functional connections of the face network and did not investigate the contribution of waveform-based cell subpopulations (Zaldivar et al., 2022).

**Figure 1.**
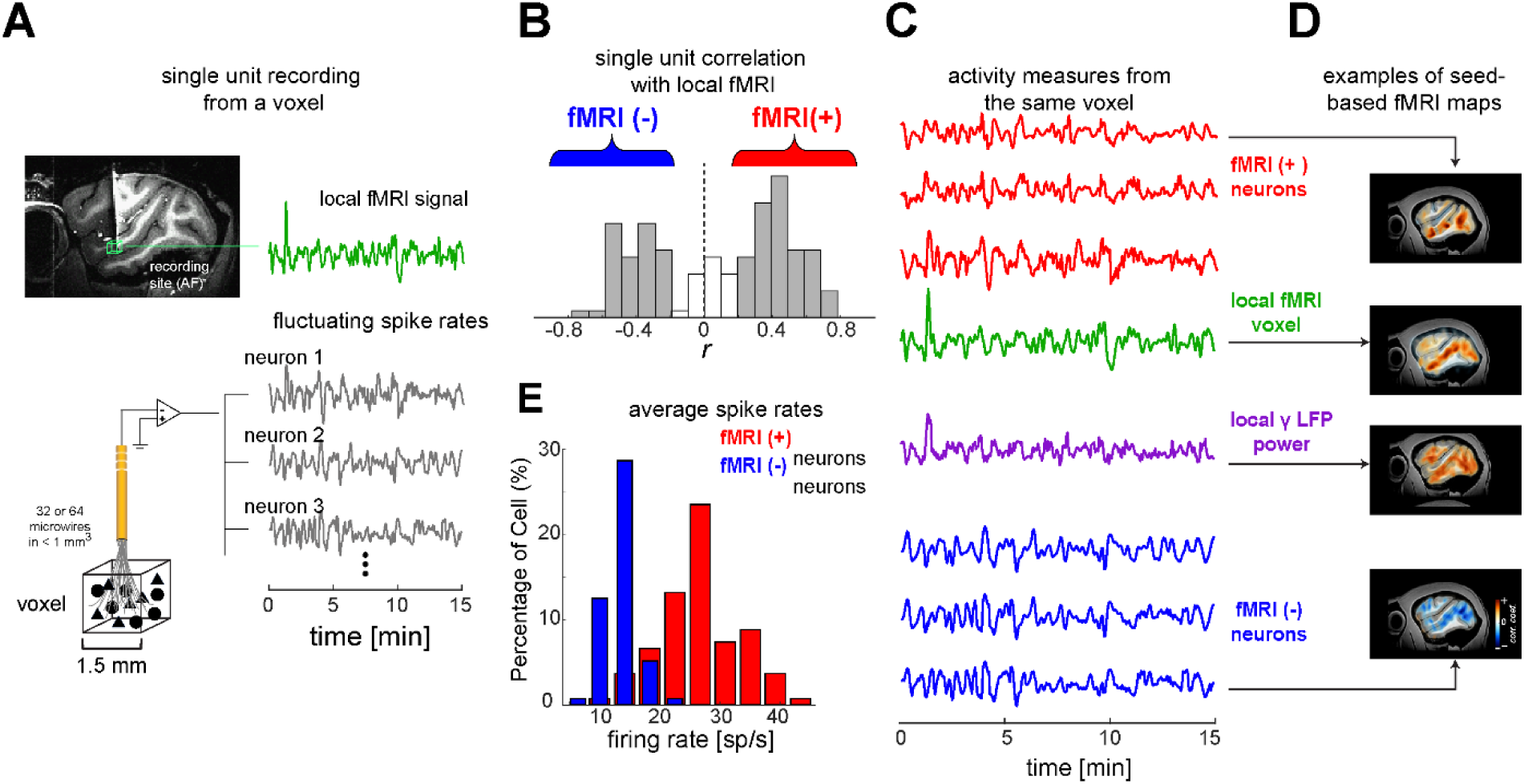
Intermixed neural subpopulations have an opposite activity relationship to the local fMRI signal. **A.** Schematic of concurrent fMRI and single-unit recordings in the resting macaque. A structural MRI example showing the location of MRI-compatible brush array electrode that was chronically and densely sampling single units within a 1.5 mm³ cortical volume. The local fMRI signal (green time-course) was measured together with fMRI signals from all voxels in the brain (not shown) along with the spontaneous activity time course of multiple isolated neurons (gray time-course). The spiking time courses were convolved and sampled to match the fMRI signal to facilitate comparison (Zaldivar et al., 2022). **B.** Histogram of correlation between local fMRI signal and all isolated single units. The distribution had clear positive and negative peaks indicating neurons that were correlated, fMRI(+), and anticorrelated, fMRI(-), with the local fMRI signal, respectively. **C.** Example time-courses from fMRI(+) neurons, fMRI voxels, fMRI(-) neurons (shown in red and blue, respectively), and gamma-band local field potential (γ-LFP) power (shown in purple). Local fMRI voxel signals positively correlate with fMRI(+) neuronal activity and γ-LFP power, while negatively correlating with fMRI(-) neuronal activity. **D.** Representative examples of seed-based functional connectivity maps derived from an fMRI(+) neuron (top), local fMRI seed and γ-LFP (middle), and fMRI(-) neuron (bottom). **E.** Histogram showing the pronounced difference in the average firing rate of fMRI(+) and fMRI(-) neurons.

### Intermixed functional cohorts correspond to waveform-based cell types

Consistent with previous findings, we observed intermixed populations of cortical neurons, the majority of which displayed spiking fluctuations that were either positively or negatively correlated with the local fMRI response (**Fig. 1B**, (Landemard et al., 2026; Liu et al., 2021; Zaldivar *et al*., 2022)). One subset of neurons (46.5%, 73/157) exhibited strong or moderate positive correlation (r > 0.2, fMRI(+)), whereas a different subset (29.9%, 47/157) exhibited strong to moderate negative correlation (r < -0.2, fMRI(-)). The remaining neurons (23%, 37/157) had weak or no correlation (|r| ≤ 0.2, S0). Accordingly, the spiking fluctuations between were positively correlated between pairs of neurons within each group (r = 0.61 ± 0.21 and r = 0.56 ± 0.17 for fMRI(+) and fMRI(-), respectively) and negatively correlated between the two groups (r = -0.56 ± 0.23) (**Fig. S2A**). The two groups were similarly divided with respect to fluctuations in the γ-range local field potential (LFP) power (**Fig. S2B**), reflecting the close relationship between the γ-LFP power and spontaneous fMRI signal (Leopold et al., 2003; Logothetis et al., 2001). Representative time courses of positively and negatively correlated single-units, the local fMRI voxel, and γ-range LFP power (**Fig 1C**) exemplify the shared and inverted temporal motifs in the simultaneously recorded spontaneous signals. Maps of seed-based correlations across the brain were rendered from each local signal type, including single neurons, local hemodynamic responses, and γ-LFP power, using the Spearman rank correlation (**Fig 1D**).

Neurons exhibiting positive local fMRI correlations had much higher average firing rate than those with negative local fMRI correlations (**Fig. 1E)**. Whereas the fMRI(+) neurons had a mean firing rate of 27.3 ± 5.1 sp/s, the fMRI(-) neurons had a mean firing rate of 11.8 ± 2.8 sp/s. Thus, the polarity of a neuron’s spontaneous rate fluctuations relative to the local fMRI signal was well predicted by its average firing rate. This difference cannot stem from lower firing neurons having effectively poorer signal-to-noise, since the magnitudes of the positive and negative correlations in the two populations were similar (**Fig. 1B**).

We next investigated whether the differences in polarity and spike rate might reflect the differential contribution of distinct cell types. To this end, we applied a standard method to classify neural subtypes based on the waveform shape of their extracellularly recorded action potentials. In the macaque, this approach is commonly used to distinguish between putative inhibitory interneurons, some of which have uniquely narrow waveforms, and putative excitatory neurons, most of which have broad waveforms (Kaufman et al., 2013; Lee et al., 2021; Snyder et al., 2016; Trainito *et al*., 2019).

We analyzed spike waveform features of all 157 neurons using a three-step strategy similar to previous studies without regard to experiment-related parameters such as the firing rate or fMRI correlation (Peyrache et al., 2012) (**Fig. 2A**). Briefly, as the first step, averaged spike waveforms from each neuron were temporally aligned and subjected to analysis of their key waveform features. These features, which included trough-to-peak time (the interval between the global minimum and subsequent local maximum), half-peak width (measured at 50% of the peak amplitude, capturing the spike width at mid-height and reflecting depolarization and repolarization kinetics), repolarization time (RT; the interval from spike peak to return toward baseline (Lee *et al*., 2021; Snyder *et al*., 2016)), and maximum rise slope (the steepest voltage change during depolarization (Barzo et al., 2025; Ma et al., 2024), were extracted and compiled into a multidimensional feature matrix as the second step. Finally, a principal component analysis (PCA) was used to optimize the dimensions in this feature space, allowing neural subgroups to be identified using k-means clustering (**Fig. 2B**).

**Figure 2.**
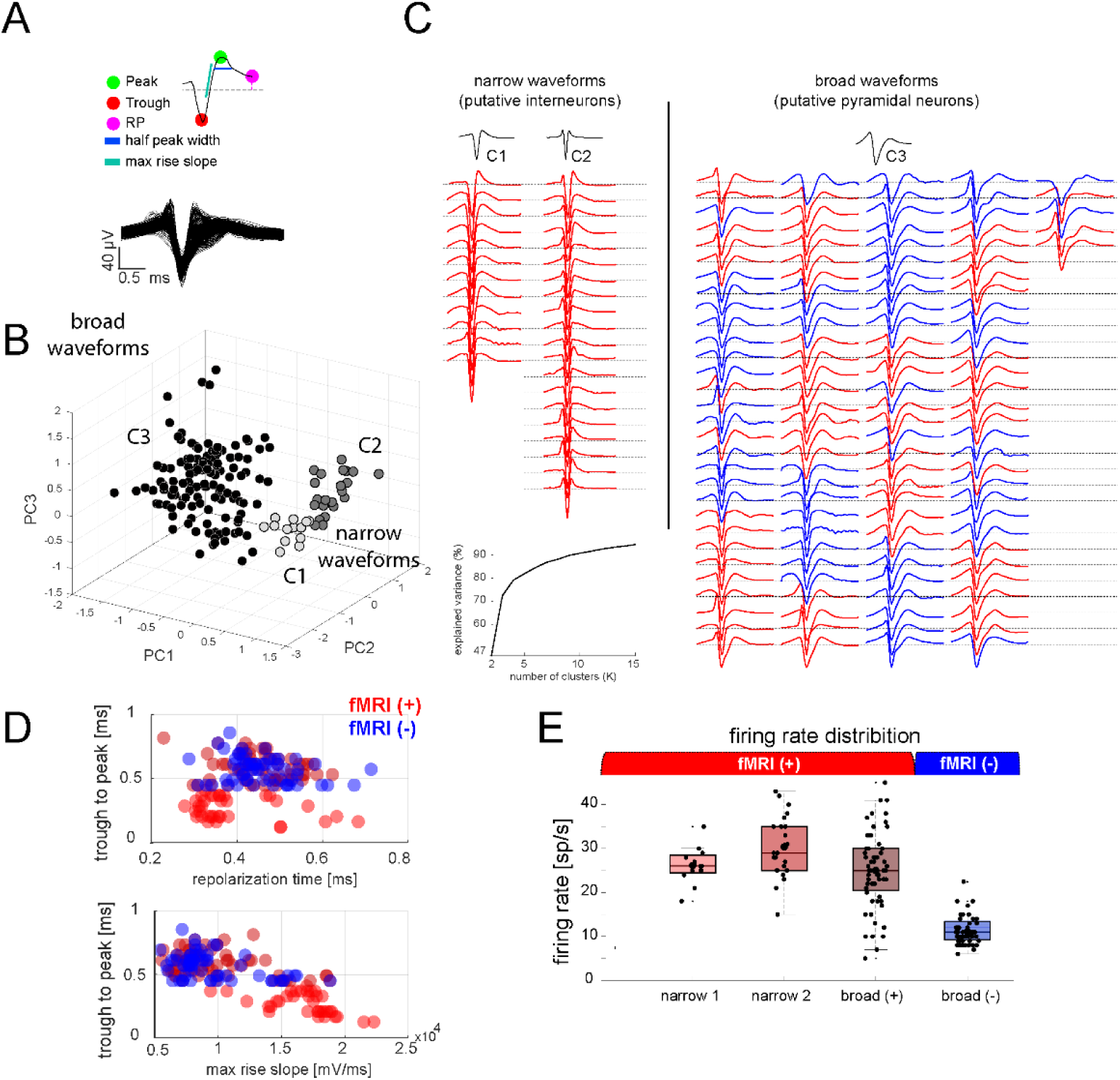
Neurons positively correlated with the fMRI signal have narrow waveforms and high firing rates. **A.** Spike waveform features—including trough-to-peak (TP) time, repolarization time (RT), half-peak width, and maximum rise slope—were extracted from all isolated single units. Overlaid average spike waveforms illustrate the diversity of waveform shapes observed in the dataset. **B.** Waveforms were subsequently classified using k-means clustering based on the extracted waveform features, revealing three distinct waveform clusters (inset represent the explained variance as a function of the number of clusters). **C.** Categorized waveforms of all 157 neurons. C1 and C2 correspond to two class of narrow waveforms defined based on spiking features, while the broad neurons represent all waveforms from the (C3) broad cluster. Red waveforms indicate neurons that positively correlated with the local fMRI signal, whereas blue neurons correspond to cells that negatively correlated to the local fMRI signal. All cell groups were sorted and plotted based on their TP duration. **D.** Pairwise scatter plots depict the distribution of neurons across waveform features: TP vs. RT (top) and TP vs. maximum rise slope (bottom). Distinct distributions of fMRI⁺ and fMRI⁻ neurons suggest systematic differences in spike waveform properties between groups. **E.** Firing rate distributions across neuron types and fMRI correlation groups. Boxplots show median and interquartile range, whiskers indicate the full data range, and black dots represent individual neurons. Narrow-spiking neurons (narrow 1 and narrow 2; fMRI⁺) exhibited the highest firing rates, broad fMRI⁺ neurons showed intermediate rates, and broad fMRI⁻ neurons displayed the lowest rates. These results indicate that neurons positively correlated with the fMRI signal tend to have narrow spike waveforms and elevated firing rates relative to negatively correlated neurons.

We identified three main waveform-based clusters with distinct properties. Neurons in two of the clusters (*C1*, 12 neurons and *C2*, 20 neurons) had narrow action potential waveforms (**Fig. 2B**, shown in red) and a third (*C3*, 125 neurons) had broad waveforms (**Fig. 2C**, shown in gray). For example, neurons in C1 displayed a trough-to-peak (TP) duration of 0.362 ± 0.024 ms and a half-width (HW) of 0.167 ± 0.021 ms, whereas C2 neurons showed shorter TP durations (0.219 ± 0.059 ms) and comparable HW values (0.150 ± 0.023 ms). In contrast, C3 neurons exhibited broader waveforms with longer TP durations (0.580 ± 0.099 ms) and slightly increased HW values (0.183 ± 0.026 ms). Additional waveform features, including repolarization time (C1 0.37 ± 0.0274 ms, C2 0.3718 ± 0.08 ms; C3 0.48 ± 0.08 ms), further contributed to the separation of these clusters (Fig. 2B).

The number of clusters was determined based on the combination of variance explained and clustering stability, with the first three clusters accounting for approximately 78% of the total variance (**Fig. 2B**, inset). The clusters were well separated in PCA space, reflecting distinct waveform characteristics across neurons. These waveform-based distinctions close to previous descriptions of intrinsic spike-shape differences among cortical cell classes, including inhibitory populations characterized using enhancer-based approaches in macaque cortical slice recordings (Furlanis et al., 2025). However, because our classification is based on extracellularly recorded action potentials, these clusters should be interpreted as putative electrophysiological classes rather than direct molecular or anatomical cell-type identities. This analysis provided the basis for analyzing the contributions of neurons from distinct putative cell classes.

We found that the relationship to the fMRI signal and the spontaneous firing characteristics differed greatly among the putative cell classes. Notably, all narrow waveform neurons (32/32 neurons, including *C1* and *C2*), exhibited positive correlation with the local fMRI signal (**Fig. 2C**, left, **Fig. 2D**). By contrast, the broad waveform (*C3*) population was divided nearly evenly into neurons that were positively and negatively correlated with the local fMRI signal. The narrow-waveform putative interneurons (C1, C2) had high average firing rates (C1 = 26.16 ± 1.248; C2 = 30.40 ± 1.531), in agreement with previous reports (Peyrache *et al*., 2012; Trainito *et al*., 2019) (**Fig. 2E**). Among the broad waveform neurons, those with negative local fMRI correlations had notably lower average firing rates (11.8 ± 2.8 sp/s) than those with positive fMRI correlations (25 ± 1.11 sp/s, **Fig. 2E**).

Given their respective association with inhibitory interneurons and pyramidal neurons, the observed functional distinction between the narrow and broad waveform neurons may reflect the roles of different cell classes during periods of spontaneous rest. We next asked whether such classification might also bear on the interpretation of fMRI functional connectivity.

### Functional connectivity closely linked to narrow waveform neurons

We assessed the contribution of different cell classes to brain-wide fMRI functional connectivity using a seed-based approach, with conventional functional connectivity derived from the local hemodynamic seed serving as a reference (**Fig. 3A**). We focused initially on correlations across the cerebral cortex, combining data across the two cortical recording sites. Broadly speaking, the spatial pattern of functional connectivity resembled the fMRI functional connectivity for all four neuron subclasses, albeit with an inverted correlation map for the negatively correlated broad population (**Fig. 3B**, top). However, details of the positively correlated cell classes differed, both in the mean (**Fig. 3B**; see **Fig. S4** for both hemispheres) and across the raw correlation maps of each individual neuron (**Fig. S3**). Comparing the two narrow waveform subpopulations, the *narrow2* group correlations were more spatially restricted than the *narrow1* group (**Figs. 3B**, **3C**). Second, the correlation patterns from both narrow waveform groups showed much higher within-group homogeneity than those of the positively correlated broad waveform neurons (**Fig. S3**).

**Figure 3.**
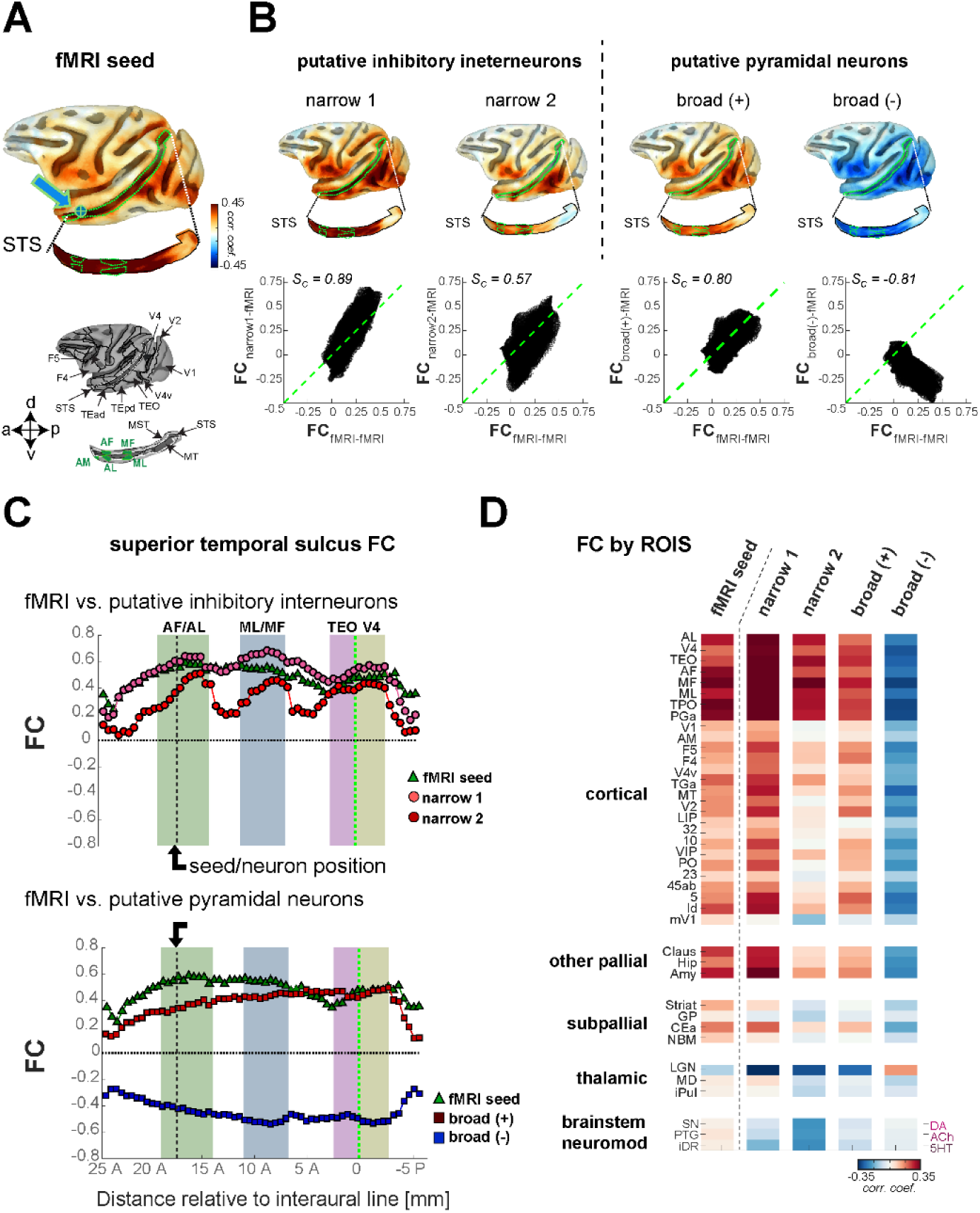
Brain-wide fMRI correlations from one subpopulation of narrow waveform neurons closely matches the fMRI seed-based functional connectivity. **A.** Lateral view of the macaque cortical surface, showing functional connectivity based on the local fMRI seed (n = 57 scans)**. B.** Lateral cortical surface maps showing functional connectivity for four waveform-defined neuronal cell types. Each map is accompanied by a voxel-wise similarity scatter plot (bottom panels) comparing the seed-based fMRI connectivity map with functional connectivity maps derived from individual neuronal groups. Cosine similarity was computed across all cortical voxels using maps obtained from the group-averaged functional connectivity patterns. Higher values indicate closer alignment with the seed-based network. **C.** Regional correlations along functionally related cortical areas defined by electrode positions in the temporal cortex. The x-axis denotes distance along the superior temporal sulcus (STS), extending from area V4 to the temporal pole. (Top) Correlation strength for the two narrow-spiking neuronal subtypes relative to the fMRI seed (green); all neurons in this group were fMRI-positive. (Bottom) Correlation strength for broad-spiking neurons relative to the fMRI seed, including both fMRI-positive and fMRI-negative populations. **D.** Region-wise functional connectivity for the fMRI seed and for group-averaged neuronal functional connectivity maps from each waveform-defined cell class. Mean functional connectivity values were extracted from 40 anatomically defined regions of interest spanning occipito-temporal visual cortex, face-selective patches, parietal and prefrontal association cortices, striatum, thalamus, and major neuromodulatory nuclei.

To establish how these differences might relate to fMRI functional connectivity, we assessed the voxel-by-voxel correspondence of neuron-based functional connectivity values (e.g. FC_narrow1-fMRI_) with corresponding fMRI functional connectivity values (FC_fMRI-fMRI_) (**Fig. 3B**, bottom). For each comparison, functional connectivity maps were vectorized by collapsing cortical data into a linear vector of 90,112 vertices. The similarity to the fMRI seed-based reference maps was quantified using similarity analysis (S_C_), which measures the degree to which the overall spatial pattern of connectivity is aligned with the reference map, independent of absolute magnitude (Bajada et al., 2020; Hong et al., 2019).

Among all neuronal groups, the *narrow1* class of putative interneurons exhibited the strongest similarity to the fMRI seed-based functional connectivity pattern (S_C_ = 0.88, *p < 5x*10^-5^, 20,000 permutations). This correspondence was reflected in a tight, elongated distribution of paired correlation coefficients between the neural and fMRI seed-based maps, with a slope that was slightly steeper than the identity line. This relationship indicating a strong and systematic correspondence of the cortex-wide functional connectivity map derived from the *narrow1* neurons with that computed from the local fMRI seed. The *narrow2* group showed a broader and more dispersed distribution, with reduced alignment to the identity line, indicating lower fidelity and increased variability compared to *narrow1*, though the relationship remained statistically significant (S_C_ = 0.57, *p <* 5×10^-5^, permutation test). Broad-waveform neurons displayed structured but distinct voxel-wise relationships relative to the same seed map: *broad (+)* neurons maintained a coherent linear relationship centered around the identity line (S_C_ = 0.80, *p < 5*x10^-5^, permutation test), whereas *broad (–)* neurons exhibited a similarly organized, yet sign-inverted, relationship across voxels (S_C_ = −0.81, *p < 5*x10^-5^, permutation test). Together, these analyses indicate that spontaneous fluctuations of *narrow1* neurons are most accurately reflected in fMRI seed-based functional connectivity.

We next assessed the regional distribution of correspondence in the vicinity of the recording sites. The anterior fundus face patch recorded in the superior temporal sulcus (STS), but virtue of its proximity to other STS face patches, provided an excellent opportunity to evaluate mesoscopic variation in functional connectivity. We plotted the functional connectivity of each cell group for positions along the STS-V4 axis (**Fig. 3C**; see **Methods**). This analysis again revealed the *narrow1* neuron group had the strongest correspondence with fMRI functional connectivity (**Fig. 3C**, upper plot). The *narrow2* correlations in the STS-V4 axis were weaker overall, with a pattern of peaks in the locations of the known face patch territories (for separate analysis of neurons from the two recording locations, see **Fig. S5**). This spatially restricted pattern, together with the waveform being the narrowest of the cells we recorded, may suggest that the narrow2 neurons correspond to the PV+ interneurons subclass (see **Discussion**). The spatial pattern of mean correlation of the *broad (+)* and *broad (-)* groups were continuous along this axis and, while inverted, roughly similar in magnitude, increasing slightly from anterior to posterior STS and then declining in area V4 (**Fig. 3C**, lower plot).

The pattern of neural correlation was generally similar between the two hemispheres, matching the fMRI functional connectivity (**Fig. S6A**). However, there was one notable exception along the STS/V4 axis in the vicinity of the AF recording location (**Fig. S6B**). While the fMRI functional connectivity was bilaterally symmetrical throughout this region, the neuron-based correlations near the recording site were significantly stronger on the side of the electrode. While such asymmetry was evident in all neural subpopulations, it was most pronounced in the narrow waveform neurons and specifically for the *narrow2* neurons exhibiting the more spatially restricted correlations. For this cell group, there the hemispheric difference in correlation at the AF recording location exceeded 0.3. This asymmetry contrasted sharply with the near absence of local hemispheric difference computed from the local fMRI seed.

We next examined seed-based correlations within fMRI fluctuations in several regions of interest (ROIs), including noncortical structures such as the thalamus, amygdala, and brainstem neuromodulatory centers (see **Methods** for ROI definitions; see **Fig. S7-S9** for whole-brain analyses). Comparing the profile of fMRI seed-based functional connectivity (FC_fMRI-fMRI_) to the average correlation value of all neurons in each class (**Fig. 3D**) revealed some similarities, such as the highest magnitude correlation in the vicinity of the STS recording sites (top eight ROIs). In other areas, and particularly subcortical structures such as the thalamus and brainstem, the functional connectivity of the different cell populations showed greater deviation from the fMRI seed-based functional connectivity.

We next investigated these ROI profiles for all 157 individual neurons across all five monkeys (**Fig. 4A**). For each neuron, we quantified the similarity of the ROI connectivity profile to the corresponding fMRI seed-based functional connectivity (**Fig. 4B**). In agreement with the cortex-based analysis above, the strongest overall correspondence with fMRI functional connectivity was evident in the *narrow1* class (S_C_ = 0.92). This population also showed the most uniform functional profile across the waveform-defined neural subtype. The ROI profiles of *narrow2* neurons were more diverse and exhibited overall less similarity to the fMRI functional connectivity profile (S_C_ = 0.62). In areas such as the parietal cortex, early visual cortex, claustrum, and amygdala, the correlations of the *narrow1* neurons showed stronger correspondence to the fMRI functional overall diversity in their ROI correlation profiles. Beyond varying in their overall polarity, this heterogeneous group included some neurons with functional connectivity largely restricted to STS locations and others with and much broader functional connectivity. The broad waveform group also contained neurons with unique connectivity patterns across specific ROI combinations.

**Figure 4.**
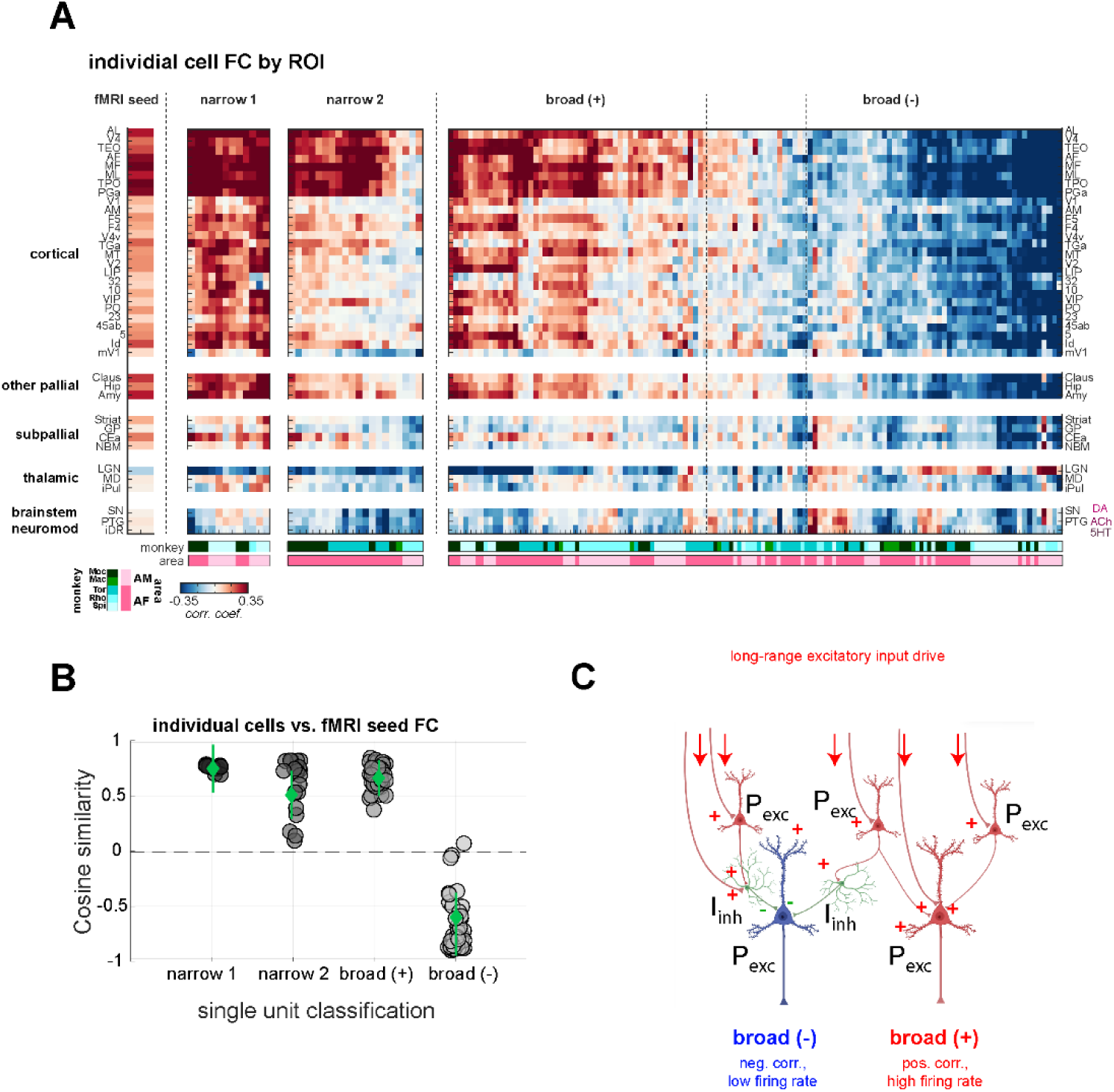
Seed-based resolved functional connectivity profiles computed from cortical and subcortical regions of interest (ROI). **A.** Region-wise functional connectivity (FC) estimated for the fMRI seed (left) and for the group-averaged neuronal FC patterns of each waveform-defined cell (right). Mean FC values were extracted across 40 anatomically defined ROIs spanning occipito-temporal visual cortex, face-patch regions, parietal and prefrontal association areas, striatum, thalamus, and major neuromodulatory nuclei. This panel illustrates that the distinct neuronal subclasses exhibit characteristic large-scale FC signatures, enabling direct comparison between the canonical fMRI seed map (left) and the aggregate FC patterns produced by each cell group (right). Neurons are ordered based on their waveform (narrow 1, narrow 2, broad (+), and broad (–)). The S0 of cells composed by broad cells can be found in between the broad (+) and broad (–). Each column corresponds to a single neuron and each row to an ROI. Color values represent the average correlation strengths. Bottom heatmaps encode the monkey and the cortical area from which each neuron was recorded, aligned to the neuron-resolved FC, suggesting that these functional distinctions are conserved across subjects and recording locations within the face-patch system. **B**. Similarities between cortical fMRI functional connectivity maps derived from individual neurons and the seed-based fMRI connectivity map, grouped according to waveform class. Narrow 1 neuron show the strongest correspondence to the seed-based map, whereas the other waveform-defined groups exhibit greater heterogeneity in their similarity to the canonical fMRI connectivity pattern. Group-level quantification of similarity values across waveform classes reveals systematic differences in how closely each neuronal subclass recapitulates the seed-based fMRI connectivity profile, indicating that waveform-defined cell types differentially contribute to large-scale functional network organization. **C.** Schematic model illustrating how the neural subclasses may differentially shape large-scale fMRI connectivity patterns. Excitatory pyramidal neurons (red) and inhibitory interneurons (green) receive direct and indirect long-range excitatory input. Downstream local excitatory neurons receiving stronger feedforward inhibition (broad (-)) exhibit inversed polarity of their fluctuations along with reduced firing rates. In contrast, excitatory neurons receiving predominantly excitatory propagation retain their polarity and have relatively higher firing rates. This model the predominance of positive polarity putative interneurons, along with the observation that even among putative excitatory neurons negative signal polarity is accompanied by decreased firing rate. According to this model, interareal excitatory projections would carry the positive polarity signal (i.e. broad (+)). Other potentially important circuit motifs, such as those involving disinhibition, are not depicted, in part because no negative polarity interneurons (i.e. narrow (-)) were observed.

As noted above for the population average responses, and as reported previously in reference to face patch connections (Zaldivar et al., 2022), neural functional connectivity deviated significantly from fMRI functional connectivity in thalamic and brainstem neuromodulatory areas. High magnitude correlations with the LGN, generally inverted in polarity relative to the cortical correlations, were prominent among both classes of putative interneurons as well as a subset of the broad waveform neurons (**Fig. 4A**). The *narrow1* and *narrow2* neurons differed in functional connectivity with other thalamic nuclei and brainstem neuromodulatory centers: in both cases, the *narrow2* neurons showed prominent negative correlation with these structures whereas the *narrow1* neurons did not.

In sum, the functional connectivity computed from different neural subpopulations differed in their relationship to one another and to traditional fMRI seed-based functional connectivity. The fMRI seed-based functional connectivity most closely matched the spatial pattern derived from one subpopulation of putative inhibitory interneurons, which all had positive correlation with the local fMRI signal. The more numerous putative excitatory projection neurons varied markedly in both their pattern of their functional connectivity, with neurons having negative local fMRI correlation. Surprisingly, this analysis suggests that the best predictor of resting-state functional connectivity across the cortex is not the fluctuating activity of projection neurons but rather that of local interneurons. Finally, for some subcortical structures, fMRI seed-based functional connectivity did not clearly match the functional connectivity derived from any of the neural subtypes.

## Discussion

We provide a comprehensive analysis of the relationship between fMRI functional connectivity and the activity of neural subpopulations within a cortical voxel. Neural classification based on waveform shape was a good predictor of a neuron’s relationship to fMRI functional connectivity.

### Neural subtype contributions to brain-wide functional connectivity

A key finding of the current study was that during rest different cell classes exhibited distinct spatial patterns of functional connectivity, as well as different levels of heterogeneity within each waveform-defined category. An unexpected result was that narrow-waveform putative local interneurons all showed positive correlation with the local fMRI signal, whereas the broad waveform neurons were roughly evenly divided in having positive and negative correlations (**Fig. 4A, Fig. S3**). This pattern suggests that fMRI functional connectivity may be more closely linked to neural activity associated with local processing than to activity specifically reflecting long-range projection output. It may also reflect a privileged role of interneurons mediating local hemodynamic responses (Elorette *et al*., 2024; Sundqvist et al., 2025; Tremblay *et al*., 2016; Uhlirova *et al*., 2016; Vo *et al*., 2023; Vo *et al*., 2025), in line with evidence that functionally distinct contributions can arise within spatially intermixed neural populations (Liu *et al*., 2021). At the same time, the coexistence of opposing correlation polarity signals within a cortical voxel, together with the uniformly positive coupling of narrow waveform neurons to the hemodynamic signal raise a key mechanistic question, namely, what is the nature of spontaneous signals transmitted through long-range projections thought to drive functional connectivity by synchronizing across brain areas? We address this question in the next section.

Among the putative interneurons, we found differences between the two waveform-defined populations. For example, *narrow2* neurons showed highly regionalized correlations compared to *narrow1* neurons. As the recording sites were in fMRI-localized face patches, this was manifest as discrete regions of correlated activity within nodes of the face patch network (**Fig. S6**), seeming to match known underlying anatomical connections (Grimaldi *et al*., 2016). The same neurons also showed the strongest hemispheric asymmetry in the fMRI correlations near the recording site (**Fig. S6**). This finding may suggest that the *narrow2* interneuron subtype is selectively or differentially driven by excitatory inputs from other face patches. Functionally selective input onto specific cortical interneuron populations has previously been demonstrated in other systems (Naskar et al., 2021). This restricted processing may serve to focus activity within specialized circuits, in this case among neurons communicating within the face network. None of the other cell classes, including the *narrow1* neurons, exhibited this level of spatial restriction. The broader correlations of *narrow1*, which included nodes of the default mode network, may have a closer relationship to neuromodulation. They may also reflect other facets of cortical inhibition, such as those related to offsetting the waves of excitation that accompany spontaneous fluctuations (Isaacson and Scanziani, 2011).

Regardless, it is interesting that the functional connectivity of the *narrow1* neurons more closely matches with the fMRI seed-based functional connectivity, rather than that of the more precisely correlated *narrow2* neurons. How these subpopulations map onto known interneuron cell types, such as PV+ and SST+ cells, is presently a matter of conjecture. Recent work suggests that these two classes of interneurons may have distinct roles in feedforward and feedback inhibition, which may link to the observed differences in spatial specificity (Jang et al., 2020). This potential distinction between these two populations draws attention to the fact that fMRI seed-based functional connectivity typically does not consider this factor (however, see (Lawrence et al., 2019). In addition, it highlights the fact that fMRI functional connectivity does not identify the most reduced network representation but is instead an inherently composite pattern driven by a superposition of physiological mechanisms.

By comparison, the broad waveform neurons recorded from the same locations were far more heterogeneous in their correlation patterns. For example, many showed promiscious functional connectivity that included cortical regions lacking well established anatomical connections. This pattern was particularly evident in the negatively correlated *broad(-)* neurons, many of which were strongly correlated with the default mode network, such as the retrosplenial, posterior cingulate, and posterior parietal cortices (Arsenault *et al*., 2018; Hutchison et al., 2011; Liu et al., 2019; Mantini et al., 2011; Turchi *et al*., 2018; Vincent *et al*., 2007; Yacoub et al., 2020). It is possible that, in addition to corticocortical connections, common-input neuromodulatory pathways or other brain-state related mechanisms play a role in shaping these correlations, as well as prominent correlations in the thalamus and elsewhere (Dearnley et al., 2023; Landemard *et al*., 2026; Liu *et al*., 2021; Lozano-Montes et al., 2020; Nair et al., 2018).

### Linking fMRI correlation, cell type, and average firing rate

The pronounced difference in firing rates between positively and negatively correlated neurons, including between the *broad(+)* and *broad(-)* categories, allows us to speculate on potential underlying mechanisms. The high firing rate among narrow waveform neurons might be expected, as some of these putative interneurons may be inherently fast spiking neurons (Henze *et al*., 2000; Huang and Paul, 2019; Milicevic et al., 2024; Trainito *et al*., 2019). The broad waveform cells are generally interpreted as putative pyramidal projection neurons, though extracellular waveform classification is not perfectly cell-type specific. Notably, the broad spiking cluster likely also contains a sampling of broad SST and VIP cells as well, which may account for some of the functional diversity (Jung et al., 2023; Onorato et al., 2025; Petersen et al., 2021; Torres-Gomez et al., 2020; Vigneswaran et al., 2011).

That being said, if adopts a simplified view that the broad waveform neurons are primarily excitatory pyramidal cells and the narrow waveform neurons are inhibitory interneurons, the link between cell class, firing rate, and correlation polarity can be captured in a simple local circuit model (**Fig. 4C**). In this model, incoming long-range excitatory projections synapse onto both pyramidal neurons and inhibitory interneurons, which in turn influence the activity of other neurons in the local circuit. These excitatory inputs would convey positive polarity fluctuations (relative to the local hemodynamic signal) that would serve to synchronize between nodes of a functional connectivity network. Locally, however, these incoming signals would variably impact excitatory neurons depending on the nature of the local interaction with inhibitory interneurons. For a subset of downstream pyramidal neurons shaped primarily by their interneuron inputs, mediating inhibition would invert the polarity of spiking fluctuations and reduce the mean firing rate (**Fig. 4C**, *broad(-)*). The inverted polarity of such cells contrasts the maintained polarity of the *broad(+)*, which are driven principally by excitatory inputs. Notably absent in this simplified model, as in our data, are interneurons carrying signals with inverted polarity, suggesting a lack of disinhibition (e.g. one interneuron inhibiting another). This absence of such neurons may either suggest that disinhibition does not play an important role in this process or brain state, or that interneurons receiving inhibition, and thus exhibiting inverted polarity, are present but classified as broad waveform neurons (Petersen *et al*., 2021).

One of the best studied interneuron subclasses is PV+ interneurons. Through the tight suppression of activity of nearby pyramidal neurons, these neurons play a role in spatially differentiated activity (Moon *et al*., 2021; Vo *et al*., 2023). As such, it is possible that the *narrow2* population maps onto PV+ neurons, which would explain the restricted patterns of functional connectivity, such as that observed in the face patch network (**Fig. 3C**). Optogenetic studies in mice have also revealed that PV+ neuron activity can induce spatially confined vasoconstriction that depends on both cortical layer and network state (Anenberg et al., 2015; Vo *et al*., 2023), suggesting that this interneuron subclass may have multiple avenues for shaping brain-wide hemodynamic correlations.

Assuming similar functional composition across the cortex, our findings suggest that fMRI functional connectivity between any two cortical areas is best approximated by the activity of local interneurons at each location, at least for seconds-long fluctuations matching fMRI time scales. These findings raise new questions about the role of long-range interconnections in eliciting synchronization across the brain. One possibility is that the *broad(+)* neurons are a critical element for interareal neural synchronization but are only secondary in determining the fMRI response. Another possibility could be that the long-range transmission is carried by *broad(-)* neurons, with synchronization achieved using inverted signals at relatively low firing rates. A third possibility is that the corticocortical long-range projections consist of a mixture of positive and negative polarity signals. While any of these are theoretically possible, the absence of putative interneurons with negative polarity correlation makes it unclear how an inverted incoming signal would undergo a local sign reversal. Recent optogenetic experiments support the outsized role of interneurons in functional connectivity, demonstrating that their manipulation causes interregional and interhemispheric changes in fMRI functional connectivity (Moon *et al*., 2025). Thus, converging evidence suggests that interneurons play important roles both in the regulation of local fMRI fluctuations and in broader patterns of functional connectivity.

### Limitations of the present study

The present study recorded single neurons during fMRI scanning of the resting macaque. The microwire array was fashioned for MR compatibility, dense sampling, and longitudinal monitoring of individual neurons. However, its geometrical structure did not allow us to determine the laminar position of the neurons we recorded. This information would be of great value, given the different interneuron composition and projection targets of pyramidal cells in upper and lower cortical layers. Previous work in isolated somatosensory cortex of the mouse has found anticorrelated activity between superficial and deep cortical layers (Kodama *et al*., 2018), suggesting that laminar positions may critically shape functional relationships. Consistent with this idea, optogenetic and electrophysiological studies have shown that activation of inhibitory interneurons can produce layer-specific modulation of neural activity, with superficial and deep neurons responding differently to inhibitory drive. Similarly, analysis of the Allen Brain dataset also identified intermixed groups of cortical neurons with anti-correlated activity with some evidence of laminar segregation alongside significant intermixing (Liu *et al*., 2021). Nonetheless, our results indicate that even without the precise laminar resolution, distinct neuronal subpopulations exert differential influence over local and network-level fMRI signals, providing important insight into the cellular basis of functional connectivity.

Our recordings were also restricted to fMRI-defined face patches in the inferior temporal cortex. Thus, the observed network organization may not be fully representative of cortex more broadly. However, it was striking that the pattern of positively and negatively correlated neurons was preserved across sessions, monkeys, and from two spatially removed face patches. Recent work in rodents has shown that neurons across diverse cortical regions exhibit positive or negative relationships to the hemodynamic signal (Landemard *et al*., 2026) suggesting that the polarity of a neuron’s coupling to the fMRI signal may be a meaningful feature across different cortical areas. At the same time, some neurons likely undergo state-dependent fluctuations in their coupling, as reported in previous studies (Shahsavarani et al., 2023) that might lead to changes in the firing rate of the neurons (Dearnley *et al*., 2023). Future work will be critical to determine whether these robust patterns reflect general principles of cortical circuitry and hemodynamic recruitment, and the particular role played by interneuron subtypes. It will also be important to assess whether the apparent dominance of interneuron-like populations in interareal fMRI coupling persists beyond the resting state, extending into more complex or naturalistic behavioral conditions.

While the spike waveform analysis performed in this study revealed functional differences between neurons of different spike characteristics, the assignment of cells to excitatory or inhibitory classes and subclasses is necessarily uncertain. Following prior conventions, we hypothesize that the two groups of narrow waveform neurons are of the parvalbumin and somatostatin types of inhibitory interneurons (Bartho et al., 2004; Cardin et al., 2010; Jung *et al*., 2023; Lee et al., 2012), but we refrained from speculating much on which corresponds to which group. The assignment of broad waveform neurons is more difficult (Petersen *et al*., 2021). It is likely that most are excitatory projection neurons, though some may be interneurons known to have broad action potentials, such as the vasoactive intestinal peptide neurons (Jung *et al*., 2023; Zaitsev et al., 2009). Future investigations incorporating histological validation will be crucial to confirm the cellular identity and spatial specificity of these neural populations, further refining our understanding of their contribution to cortical network dynamics.

Finally, the current study did not investigate neurovascular coupling directly or through pharmacological manipulation, which limits our ability to move beyond correlation to attribute fMRI signals to specific vascular or metabolic mechanisms. Future studies employing optical imaging or localized measurements of blood flow and oxygenation could complement these findings and further elucidate the relationship between neuronal activity and hemodynamic responses.

## Conclusions

This study demonstrates that the neurons comprising a cortical voxel exhibit primarily positive and negative correlative relationships to the hemodynamic fMRI signal at rest. Those neurons most clearly identifiable as inhibitory interneurons based on their narrow action potential waveforms all show positive correlations, whereas those with broader waveforms are evenly divided between positive and negative. The narrow waveform neurons match both the polarity of local fMRI responses and the spatial pattern of longer-range pattern of fMRI seed-based functional connectivity, indicating that this neural subpopulation, more than any other, drives the signals observed in studies of resting state functional connectivity.

## STAR Methods

### Subjects and Ethical Statement

The current study represents a re-analysis of fMRI and neurophysiology data that we concurrently acquired in macaques (Zaldivar *et al*., 2022). We used six male rhesus monkeys (Macaca mulatta) weighing 7–9 kg. All animals were surgically implanted with a custom-designed MR-compatible head post and MR-compatible chronic electrodes. In five monkeys we used microwire electrode bundles to target face patches. Surgical procedures were conducted under general anesthesia using 1.5% isoflurane and were approved by the Animal Care and Use Committee of the US National Institute of Mental Health/National Institutes of Health. Following the completion of the surgery, animals were given analgesics and prophylactic antibiotics. During experiments, the animals were on water control and received their daily fluid intake during their testing. Each subject’s weight and hydration level were monitored closely and maintained throughout the experimental testing phases. All the experimental procedures were in full compliance with the Guidelines for the Care and Use of Laboratory Animals by US National Institutes of Health.

### MRI Scanning

Structural and functional MRI experiments were acquired on a vertical 4.7-T Bruker BioSpec scanner with 60-cm-diameter bore magnet (Bruker BioSpin GmbH, Germany). The Bruker S380 gradient coil with a maximal slew rate of 340 mT/m/s and a maximal strength of 56 mT/m was used. Animals sat upright in a specially designed chair. The chair base was equipped with a module housing an electromagnetic interference filter (see Neuronal Recordings section). Whole-brain images were collected with either an eight-channel receive radiofrequency coil system (RAPID MR International, Columbus, OH) or a custom-made four channel coil. A total of 67 simultaneous fMRI-electrophysiology sessions (39 sessions in AF, 28 sessions in AM) were collected. Most of the experimental sessions consisted of 600 volumes (∼30 min long), however 4 sessions in AF consisted of ∼1200 volumes (∼60 min long). Each fMRI experiment session was acquired with a single-shot gradient echo planar imaging (EPI), slice thickness of 1.5 mm, an in-plane resolution of 1.5 × 1.5 mm^2^, and a matrix of 44 × 64 × 32. Echo time and TR were 12 ms and 2,000 ms, respectively. All functional volumes in our experiments were acquired at the beginning of each TR. This was essential for our electrophysiological experiments as the gap between volume acquisition provided us with 1.2 s time window of MRI artifact-free signal. Anatomical MRI images were acquired along the same orientations as the functional images.

### MION

Prior to the start of EPI data acquisition, we intravenously administered MION. MION is a T2* contrast agent that isolates functionally related changes in cerebral blood volume, primarily in the arterioles. MION was used because of its high contrast-to-noise ratio and is common practice in monkey fMRI (Leite et al., 2002; Smirnakis et al., 2007). We obtained MION from the Imaging Probe Development Center, National Heart Lung and Blood Institute, Bethesda, Maryland. We determined the dosage of MION by monitoring the drop in intensity in the brain following injection. Previous work has shown that a drop of roughly 50% is optimal for good functional imaging, and this corresponded to roughly 8–10 mg/kg MION, depending on subject weight and batch of MION. Note that, in contrast to the BOLD signal, activity-based increases using fMRI MION signals (also known as regional cerebral blood volume) are visible as decreases in signal intensity. For the sake of clarity, we therefore inverted the sign of modulation throughout the article.

### fMRI Data Processing

AFNI/SUMA software package (Cox, 1996) and custom-written bash and MATLAB code (MathWorks, Natick, MA) were used to analyze fMRI data. Raw images were converted to the AFNI (BRIK/HEAD) format followed by slice timing correction performed in the z+ direction using the AFNI function 3dTshift with the quint (fifth order) Lagrange polynomial interpolation. Motion correction was applied to each EPI time course using the AFNI function 3dvolreg and magnetic field distortions were corrected with the PLACE algorithm (Xiang and Ye, 2007). Each scan was converted into the percentage signal change by subtracting the mean and then dividing by the mean. Global signal was not regressed out from the data due to its propensity to introduce extra negative correlations (Murphy et al., 2009) and the risk of removing physiologically meaningful signal(Schölvinck *et al*., 2010). Data from each session were aligned to a template session to facilitate the combination of data across multiple testing days (Seidlitz et al., 2018). For each monkey, spherical seed regions (2 mm diameter) were individually selected based on the corresponding recording face patch in the T1-weighted anatomical image. Fig. S1A provides examples of these regions from two monkeys. The mean time course of the signal from each seed region, subject, and scanning session was extracted and used as a regressor to perform brain-wide correlation analysis. The resulting fMRI map was then used for subsequent analysis.

### Neuronal Recordings and Analysis

Each experimental day, the setup for the simultaneous recordings involved restraining the head of the monkey and plugging the amplifier cable into the implanted connector, a procedure requiring just a few minutes. The monkey chair was equipped with an MRI-compatible electromagnetic interference filter (high-performance D-sub filter, APITech’sSeries700 EMI) which helps to reduce electromagnetic interference. The electrodes recording AF and AM face patch consisted of bundles of 32 NiCr or 64 NiCr microwires chronically implanted (Bondar *et al*., 2009). The microwire electrodes were designed and initially constructed by Dr. Igor Bondar (Institute of Higher Nervous Activity and Neurophysiology, Moscow, Russia) and subsequently manufactured commercially (Microprobe for Life Science, Gaithersburg, MD). The ground and reference were connected to one another and to a designated reference wire within the microwire bundle, as well as to a gold-plated grounding terminal adjacent to the dura, using a ceramic screw placed ∼1 cm from the implant margin. The signals obtained inside the 4.7-T scanner were amplified and digitized at 24.4 kHz with a PZ5 NeuroDigitizer (TDT) and then sorted to an RS4 Data Streamer controlled by an RZ2 BioAmp Processor (TDT). The TDT system has a resolution of ∼25 kHz and 250 nV/bit with an input range of ±500 mV, which allowed us to capture the entire electrophysiological signal with no saturation induced by the MRI artifact. Only segments of electrophysiological activity recorded during the 1.2-s gap between volume acquisitions were considered for the analysis. However, it is worth mentioning that even during these periods of MRI artifact-free signals, large residuals were still noticeable in our recordings. Therefore, we needed to account (Saggar et al., 2018; Sporns et al., 2021) for the residuals by capturing the differences from the averaged waveform. For this purpose, we used PCA as described in previous studies combining fMRI and electrophysiology (Cruttenden et al., 2022; Niazy et al., 2005). The broadband electrophysiological responses were used to extract individual spikes in WaveClus software (Quiroga et al., 2004) after filtering between 300 and 5,000 Hz. LFPs were extracted from the broadband signal by fourth-order Butterworth bandpass filtering of the signals between 1 and 150 Hz with a sharp transition bandwidth (1 Hz). Forward and backward filtering were used to eliminate phase shifts.

We classified neuron types based on electrophysiological properties derived from spike waveform shape. For each isolated neuron, we analyzed averaged spike waveforms and extracted a set of features that have been previously shown to differentiate putative excitatory and inhibitory neurons (Henze *et al*., 2000; Trainito *et al*., 2019). These features were selected independently of experiment-related variables such as firing rate or fMRI correlation.

Trough-to-peak (TP) time was defined as the interval between the waveform’s trough (global minimum voltage) and the subsequent local maximum. This measure reflects action potential duration and is known to vary systematically across neuron types due to differences in cellular morphology and ion channel composition (Connors and Gutnick, 1990; Henze *et al*., 2000). Half-peak width (HPW) was computed as the temporal width of the spike at 50% of the peak amplitude, measured from the trough of the waveform to the points on either side of the peak that reached half-maximal amplitude (Ma *et al*., 2024). This feature captures the mid-height width of the action potential and reflects the combined kinetics of spike depolarization and repolarization, with broader half-peak widths typically associated with excitatory neurons and narrower widths with inhibitory, fast-spiking neurons. Repolarization time (RT) was defined as the interval between the spike peak and the point at which the waveform returned toward baseline during the falling phase. This metric reflects the kinetics of ion channels involved in spike recovery and provides additional information about cell-type–specific repolarization dynamics. Finally, the maximum rise slope was computed as the steepest voltage change during the depolarization phase of the spike, quantifying the maximal rate of voltage increase (Barzo *et al*., 2025; Ma *et al*., 2024). This feature captures the speed of spike onset and is influenced by sodium channel dynamics, which differ across neuronal subtypes. Together, these waveform features capture complementary aspects of spike shape and dynamics, enabling systematic differentiation of neuronal subtypes based on their characteristic electrophysiological signatures.

To classify neurons quantitatively, k-means clustering was applied to the multidimensional feature matrix (TP, HPW, RT, and maximum rise slope). Because k-means is non-deterministic, the procedure was repeated 100 times for each K value from 1 to 15. For each repetition, the percentage of variance explained (between-clusters sum-of-squares relative to total sum-of-squares) was computed. Explained variance increased with K but showed diminishing returns, with no sharp inflection. Clustering stability was assessed using a co-assignment probability matrix, reflecting how consistently pairs of neurons were assigned to the same cluster across repetitions. Considering both explained variance and cluster stability, K = 3 was selected. Cluster assignments were visualized in PCA-reduced space, with distinct colors for each cluster to facilitate comparison of waveform properties across neuronal subtypes.

### Convolution of with neural spiking activity and correlation with fMRI time courses

To compare spiking activity with fMRI signals across the brain, we applied the approach used by Zaldivar et al (2022) and following earlier descriptions (Schölvinck *et al*., 2010). Briefly, we down-sampled the time course of each single unit to match the fMRI temporal resolution by taking the spike count in bins of 1.2 s, which corresponds to the 1.2 s gap between volume acquisition described above. The resulting firing-rate time series were convolved with a generic MION hemodynamic response function modeled as a gamma probability density function (shape = 3, scale = 2; (Buxton et al., 2004)) thereby accounting for the temporal characteristics of the hemodynamic response. Both neural and fMRI time courses were band-pass filtered (0.008–0.08 Hz). Following these two pre-processing steps in the time courses of each single unit, we computed Spearman’s rank correlation coefficient between the neural time course and fMRI time courses of all the voxels in the entire brain. This procedure was applied to every single unit.

### Global functional connectivity similarity analysis

To quantify similarities between the global functional connectivity maps derived from the fMRI seed FC and the connectivity maps extracted from each cell group, all functional maps were first collapsed into linear vectors of 90,112 voxels, representing the full spatial extent of all voxels across the brain. Each vector contained the voxel-wise correlation coefficient values of a single map and served as the fundamental unit for all subsequent comparisons. This approach allowed us to systematically compare the spatial patterns of FC between the fMRI-seed and each neuronal population while maintaining the voxel-level resolution.

To assess voxel-wise correspondence between the seed map and the maps from each cell group, we computed the cosine similarity. For each voxel, the similarity between the seed and the corresponding voxels in the cell group map was calculated as:

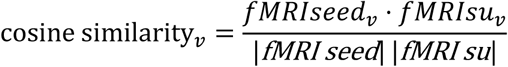

where fMRI seed and fMRI su are the vectorized functional maps, and ∥⋅∥ denotes the Euclidean norm. After computing voxel-wise and whole-map similarity metrics, we visualized the relationship between the seed map and each target map using scatter plots. For each voxel *v*, the intensity in the fMRI seed map was plotted against the corresponding intensity in the target map. This allows a direct assessment of the linearity and range of voxel-wise correspondence. To assess the statistical significance of the observed cosine similarities, we performed a two-sided permutation test for each map. Specifically, for each cell group map, the voxel labels were randomly shuffled 20,000 times, and the cosine similarity between the seed map and the shuffled map was recalculated for each permutation. The *p*-value was then defined as the proportion of permuted similarities exceeding the observed similarity. This approach allows us to determine whether the observed correspondence between the seed map and each cell group map is greater than expected by chance, accounting for the large number of voxels in each map.

We assessed similarities between the functional connectivity (FC) of the fMRI seed region and the single-unit FC extracted from each individual cell. To achieve this, we computed whole-brain weighted similarity metrics, comparing the FC from the seed (used as the reference map) to the FC from each individual group. This analysis utilized multiple approaches which have been used to compared differences in functional connectivity (Cutts et al., 2023; Hutchison et al., 2013a; Sporns *et al*., 2021), based on cosine similarity.

In addition to the global similarity analysis, we conducted a regional similarity analysis consistent of systematically sampling along a defined anatomical trajectory across the well-characterized face patch network. Specifically, we focused on activity along an STS–V4 axis, running approximately anterior-to-posterior along the superior temporal sulcus (STS) and extending onto the prelunate gyrus (Russ et al., 2023; Zaldivar *et al*., 2022). This let us generate a series of spatially ordered regions of interest (ROIs) distributed along this axis at 0.5 mm intervals, corresponding to successive coronal slices. For each ROI and functional maps from each cell type correlation values were extracted slice-by-slice from the processed fMRI maps (Fig. 3C and Fig. S5). Within each ROI, mean correlation coefficients were computed, yielding distance-resolved FC profiles along the STS–V4 trajectory for each dataset and across the different cell types. These profiles allowed us to quantify how connectivity strength evolved as a function of anatomical position, enabling direct comparison of spatial gradients across neuronal populations and between single-unit–derived and fMRI-derived FC.

### Region of Interest (ROI) analysis

To summarize the correlation profile of each neuron with cortical regions, we delineated a set of 40 ROIs based on vertices exhibiting a correlation magnitude of at least 0.2 in the grand average. We began with the D99 atlas ROI parcellations, which were subsequently refined based on the functional properties of each area and by reference to macaque connectivity maps provided by CoCoMac (cocomac.org). Boundaries between regions were further corroborated using the Paxinos macaque atlas. The resulting ROIs were then interactively adapted into the NMT macaque atlas and categorized according to standard developmental subdivisions.

### Hemispheric bias

We quantified whether functional connectivity within each group was symmetric across the two hemispheres by computing the voxel-wise correlation between the left (ipsilateral) and right (contralateral) hemisphere counterparts. To ensure proper voxel-to-voxel correspondence, the right hemisphere maps were flipped prior to analysis. All analyses were restricted to gray matter voxels, with supplementary analyses performed on the whole brain. For each map, the left and flipped-right volumes were reshaped into vectors, masked to include only voxels exceeding a minimal signal threshold, and paired voxel-wise. Pearson’s correlation coefficient was then calculated between these vectors to capture the linear similarity in the spatial organization of the maps. This voxel-wise Pearson coefficient reflects the extent to which higher-intensity voxels in one hemisphere correspond to higher-intensity voxels in the contralateral hemisphere, independent of absolute signal magnitude. To visualize these relationships, scatterplots of left-versus right-hemisphere voxel values were generated for each map pair, with each plot including the identity line (y=xy = xy=x) representing perfect symmetry.

To complement the global voxel-wise symmetry analysis, we quantified regional hemispheric asymmetries along the rostro-caudal axis of the temporal cortex using a Hemispheric Difference Index (HDI). For each ROI and neuronal group, FC values from the left and right hemispheres were extracted and normalized by the number of voxels per region. The HDI was calculated as the difference between left- and right-hemisphere values divided by their sum, such that positive values indicate stronger connectivity in the left hemisphere, negative values indicate stronger connectivity in the right hemisphere, and values near zero indicate bilateral symmetry. This approach allowed us to capture localized asymmetries at the regional level that may be obscured by global voxel-wise correlations. HDI profiles were visualized as heatmaps and bubble plots, with bubble size proportional to the magnitude of asymmetry and color encoding its direction, highlighting left-dominant, right-dominant, or symmetric regions.

## Acknowledgments

Thanks to David Yu, Charles Zhu, Katy Smith, David Ide, George Dold, and Sean Kearney for their invaluable technical assistance. Many thanks also to Drs. Soohyun Lee, Peter Bandettini, and Harsh Deshpande for helpful comments and suggestions on an earlier version of the manuscript. Functional and anatomical MRI scanning was conducted in the Neurophysiology Imaging Facility Core (NIMH, NINDS, NEI). Computational analyses utilized resources from the NIH HPC Biowulf cluster (https://hpc.nih.gov). MION was generously provided by the Imaging Probe Development Center, National Heart, Lung, and Blood Institute. This research was supported by the Intramural Research Program of the National Institute of Mental Health (grants ZIAMH002838, ZIAMH002898, and ZIAMH002899) to D.A.L., and by a DFG grant (ZA9901/1-) to D.Z. The contributions of the NIH author(s) are considered Works of the United States Government. The findings and conclusions presented in this paper are those of the author(s) and do not necessarily reflect the views of the NIH or the U.S. Department of Health and Human Services.

## Citation Diversity Statement

Recent research across scientific disciplines has highlighted a systemic bias in citation practices, where work by women and scholars from underrepresented minority groups is often under cited relative to the number of papers authored by them in the field. We acknowledge this bias and have actively sought to reference studies by women and underrepresented minority scholars wherever possible in our work.

## Author Contributions

## Supplemental Material

**Figure S1.**
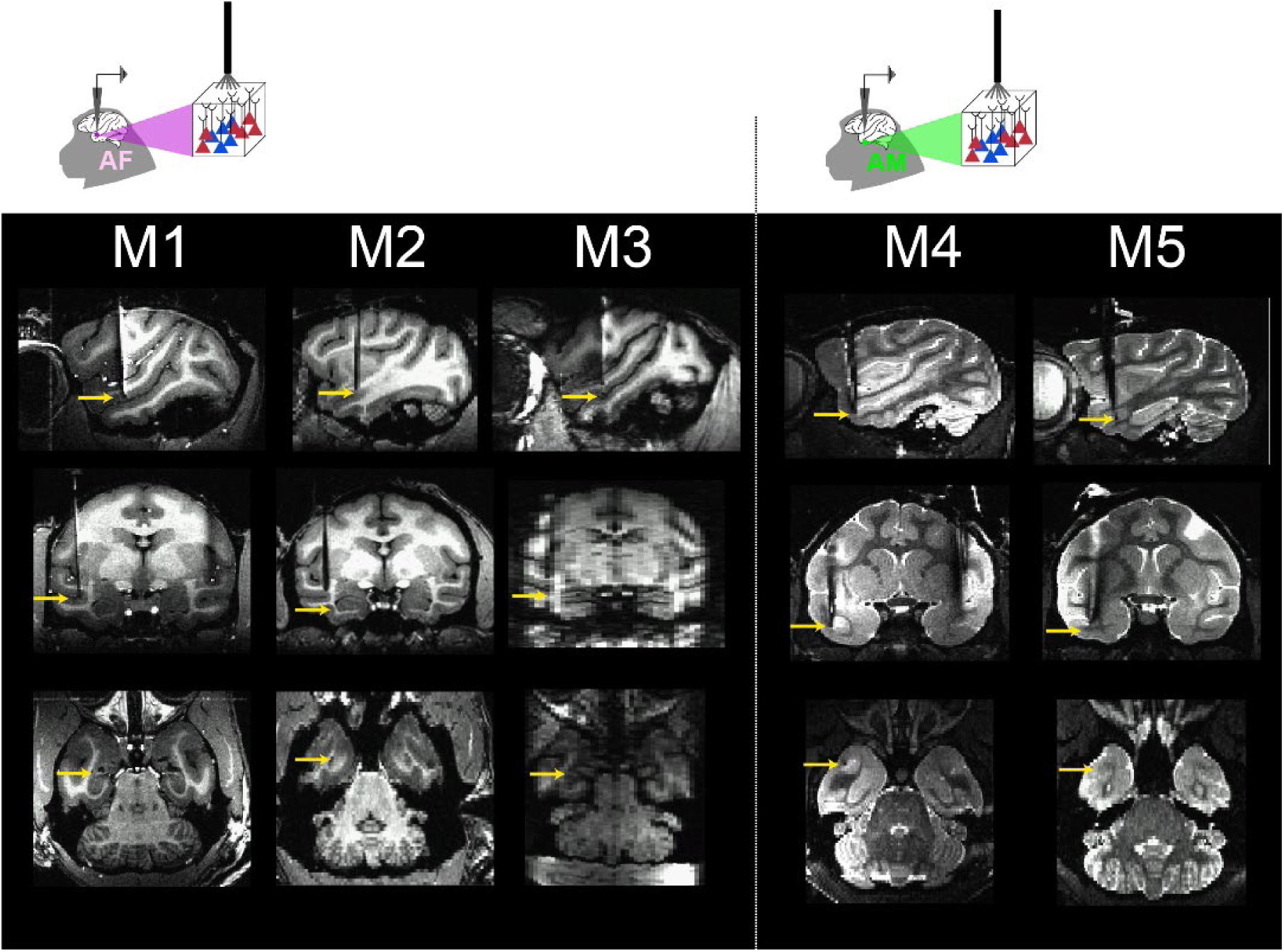
Targeting of functionally defined recording areas. Anatomical scans showing location of neurophysiological recordings in five subjects. Recordings were performed using chronically implanted 32 or 64 MR-compatible microwires. Subjects M1, M2, and M3 were implanted with electrodes in the anterior fundus face patch (left-side images) while M4 and M5 were implanted with electrodes in the anterior medial face patch.

**Figure S2.**
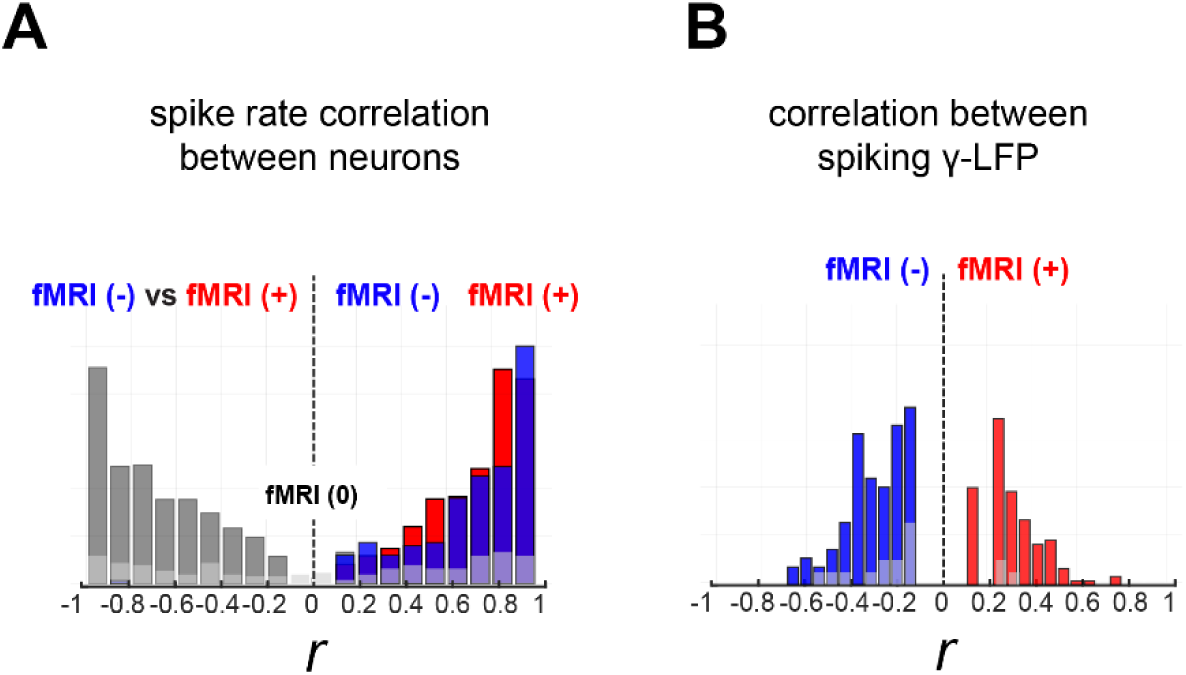
Interneuronal and spike-LFP correlations during resting state. **(A)** Distribution of spike-rate correlations between all simultaneously recorded neuron pairs. Pairwise correlations were computed either within each functional class (fMRI(+)/fMRI(+) in red; fMRI(–)/fMRI(–) in blue) or **across** classes (fMRI(+)/fMRI(–) in gray). Neuron pairs within the same class—both fMRI(+) and fMRI(–)—exhibit predominantly positive interneuronal correlations spanning a broad range of strengths, indicating coherent shared fluctuations within each population. In contrast, mixed fMRI(+)/fMRI(–) pairs cluster tightly around zero, consistent with a functional dissociation and reduced shared variability between the two classes. **(B)** Distribution of correlations between single-unit spiking activity and γ-band (40–100 Hz) LFP power. fMRI(–) neurons (blue) show strong negative correlations with γ-LFP power, whereas fMRI(+) neurons (red) exhibit positive correlations with γ-LFP power. This bivalent relationship suggests that the two neuronal classes participate in distinct local circuit states during rest, with fMRI(–) neurons preferentially associated with reductions in γ-band activity and fMRI(+) neurons associated with increases in γ-band activity.

**Figure S3.**
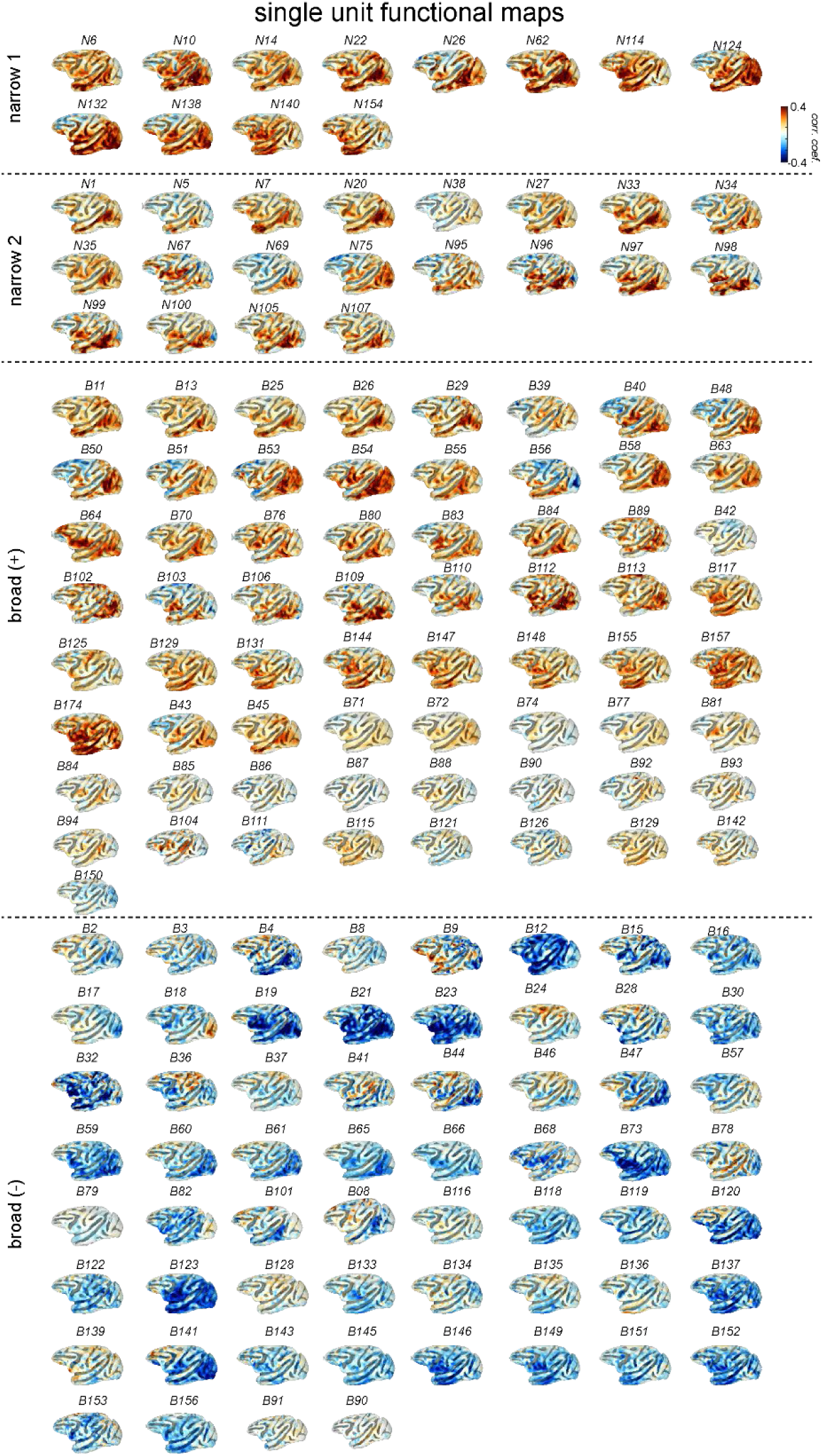
Single-unit functional maps from recorded signals in face patches. Each surface map represents single-unit functional map extracted from recordings across five monkeys. The maps are separated based on their cortical correlation map polarity and their cell type classification.

**Figure S4.**
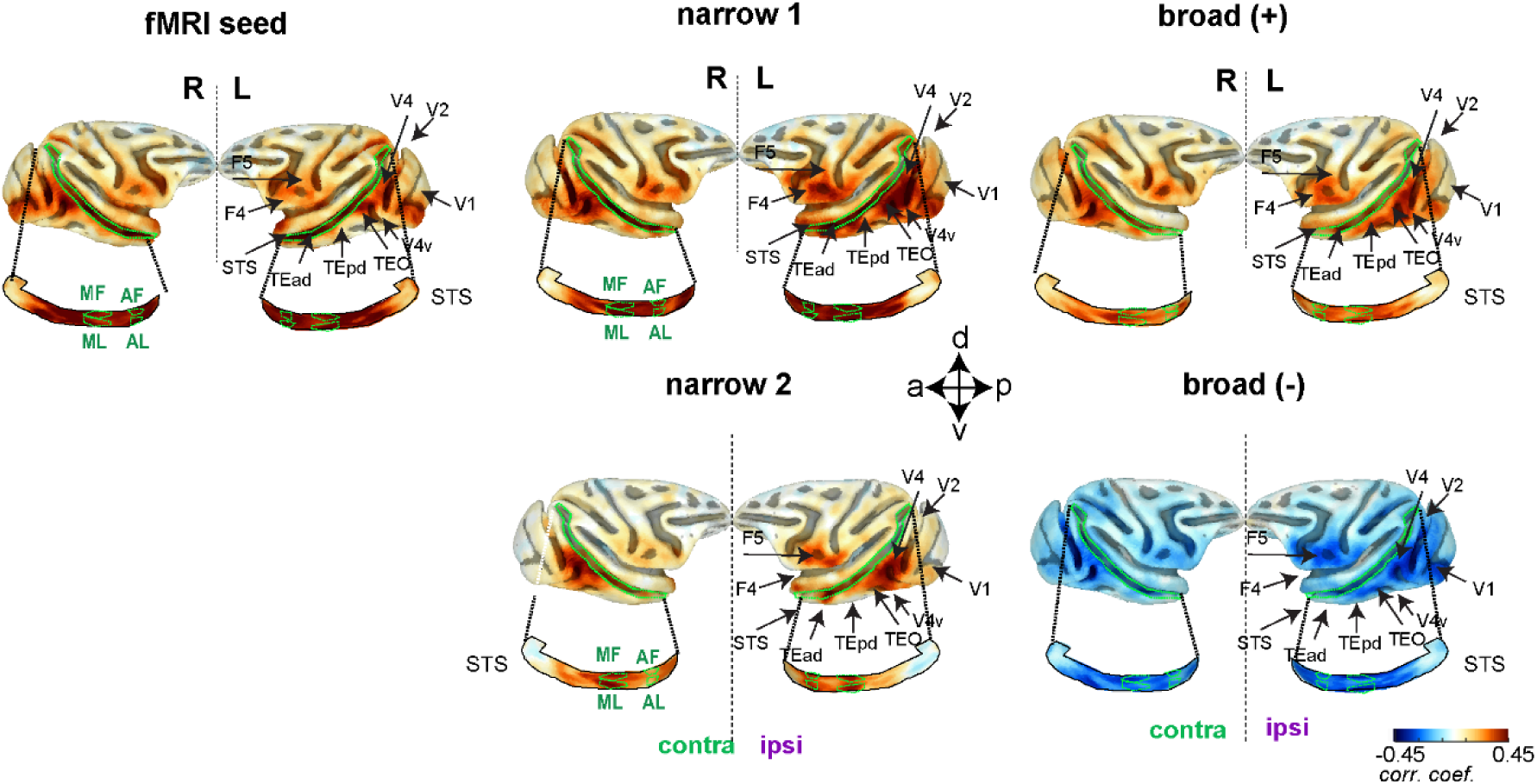
Bilateral functional connectivity for the different functional seeds. Lateral view of the macaque cortical surface showing functional connectivity (FC) for the fMRI seed and the four cell-types–defined groups. The Narrow groups (narrow 1 and narrow 2; top and bottom, respectively) consist exclusively of fMRI (+) neurons, whereas the Broad groups (broad (+) and broad (–); top and bottom, respectively) include both fMRI(+) and fMRI (–) neuron types. Boundaries of functionally defined face patches (green) are superimposed in the unfolded superior temporal sulcus (STS). Face-patch abbreviations: AL, anterior lateral; MF, middle fundus; ML, middle lateral.

**Figure S5.**
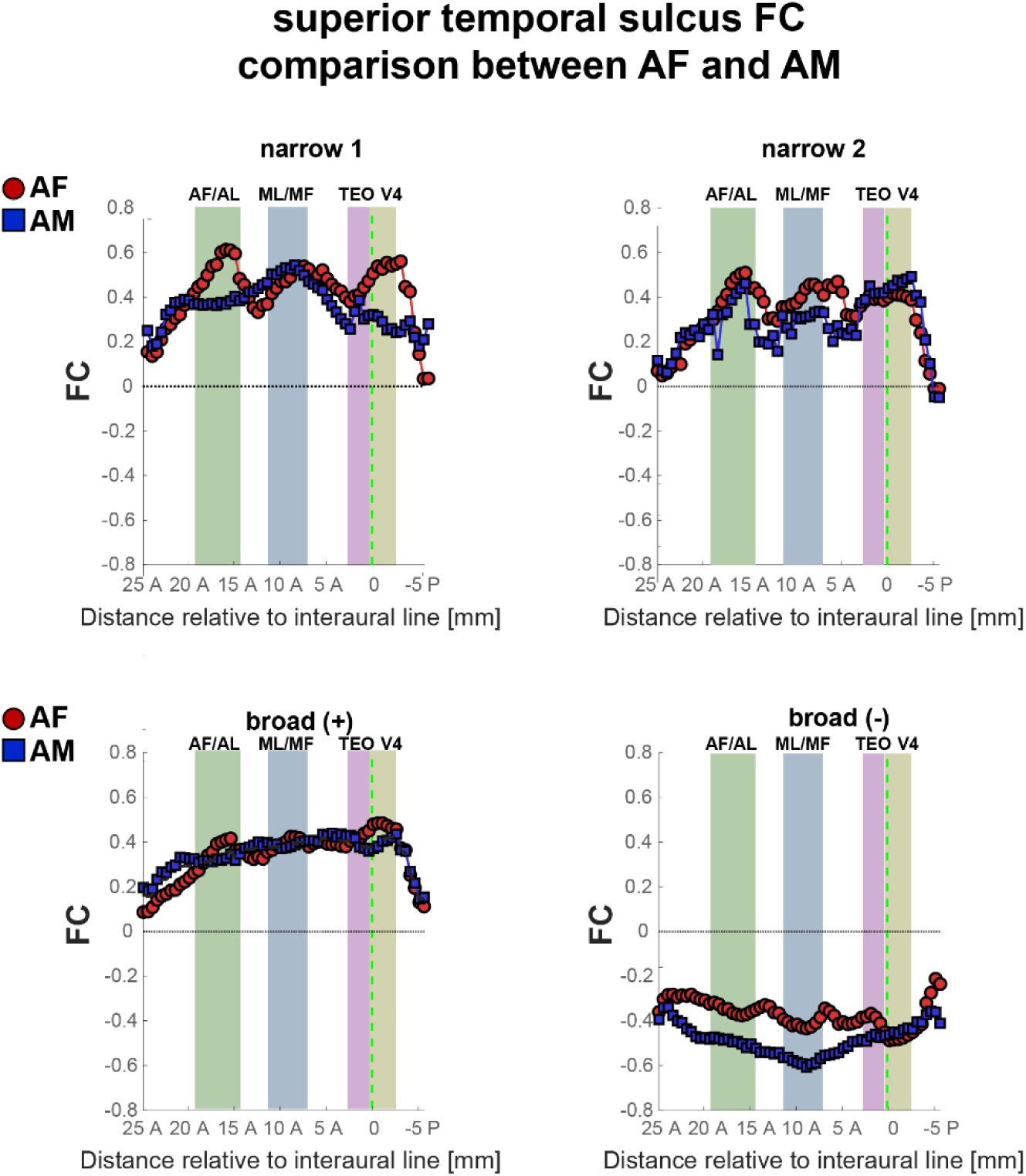
Superior temporal sulcus functional connectivity profiles for AF and AM recorded cells. Regional functional connectivity (FC) along the superior temporal sulcus (STS) is shown for cells extracted from AF face patches (red) and AM face patches (blue). The x-axis represents distance relative to the interaural line, spanning from posterior visual areas (V4) to more anterior temporal regions, including TEO, ML/MF, and AF/AL. Shaded bands indicate approximate anatomical boundaries of these regions. Data are shown separately for narrow face patches (narrow 1 and narrow 2; top row) and broad face patches with positive and negative connectivity profiles (broad (+) and broad (–); bottom row). Across conditions, AF and AM cells exhibit similar large-scale STS connectivity gradients, with systematic differences in magnitude and sign depending on patch type.

**Figure S6.**
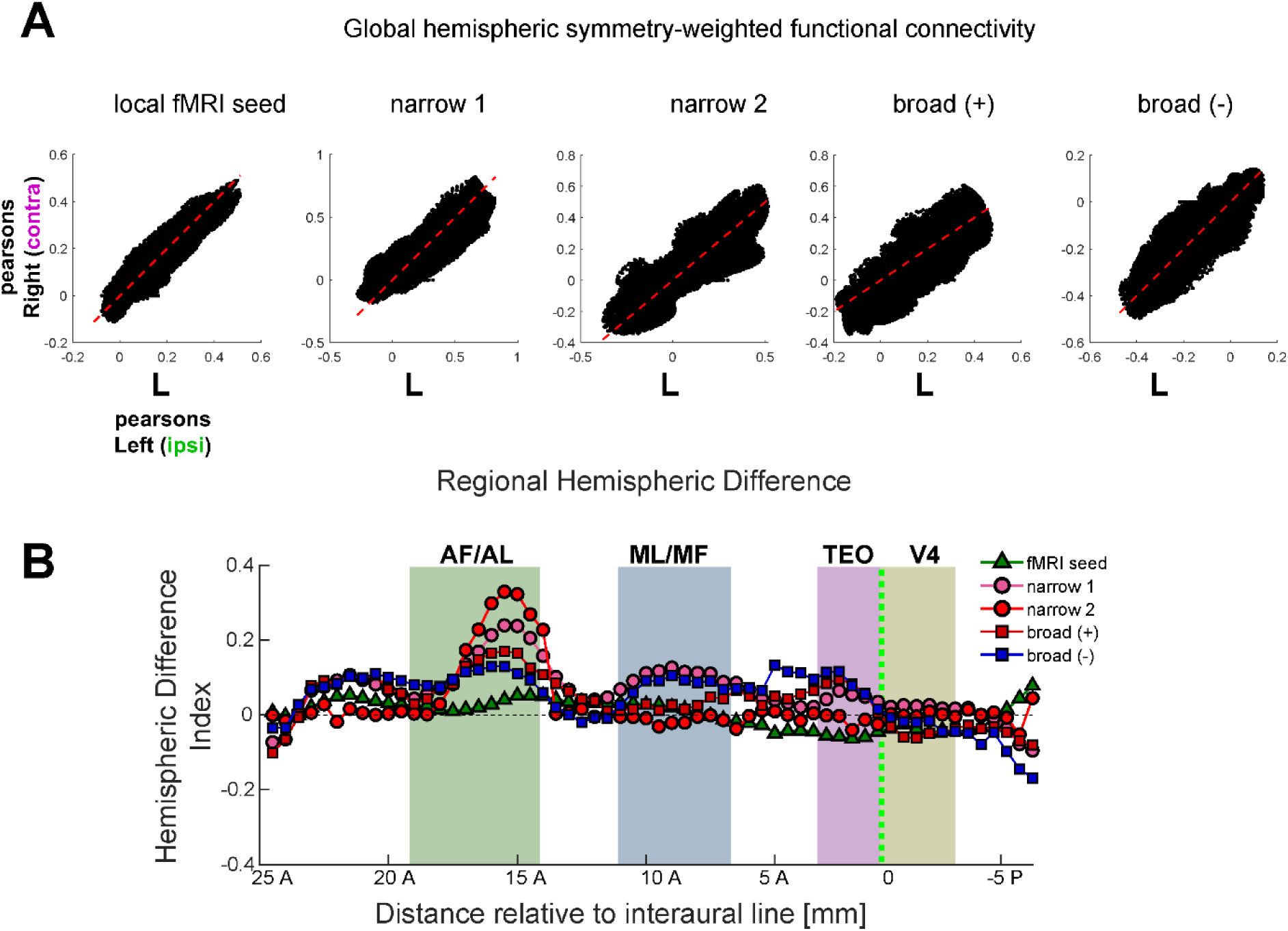
Global hemispheric symmetry of functional connectivity across neuron types. **A.** Interhemispheric correlations depict the overall connectivity patterns across different sets of functional seeds, including the local fMRI seed and seeds derived from distinct neuronal groups. The analysis shows the Pearson correlation of functional connectivity values between the left (ipsilateral) and right (contralateral) hemispheres for each neuronal group and the local fMRI seed. Each data point in the plots represents a spatially distributed voxel. The dashed red line indicates the unity line (L = R), highlighting the correspondence between hemispheres, while the blue line represents a weighted linear regression through the origin. The first panel displays correlations derived from the local fMRI seed, and the remaining panels show correlations for neuronal groups narrow 1 and narrow 2 (narrow-waveform neurons) and broad (+) and broad (–) (broad-waveform neurons). Across all groups, the functional connectivity demonstrates strong global symmetry between hemispheres. **B.** Regional hemispheric differences in functional connectivity along the STS axis. For each neuronal group and the local fMRI seed, we computed a hemispheric difference index (HDI) across the rostro-caudal axis of the superior temporal sulcus (STS). HDI was calculated as (L-R)/(L+R), where positive values indicate stronger connectivity in the left hemisphere (ipsilateral) and negative values indicate stronger connectivity in the right (contralateral). The resulting profiles show how lateralization varies regionally along the IT/V4 trajectory, revealing focal deviations from global symmetry that are specific to neuronal groups.

**Figure S7.**
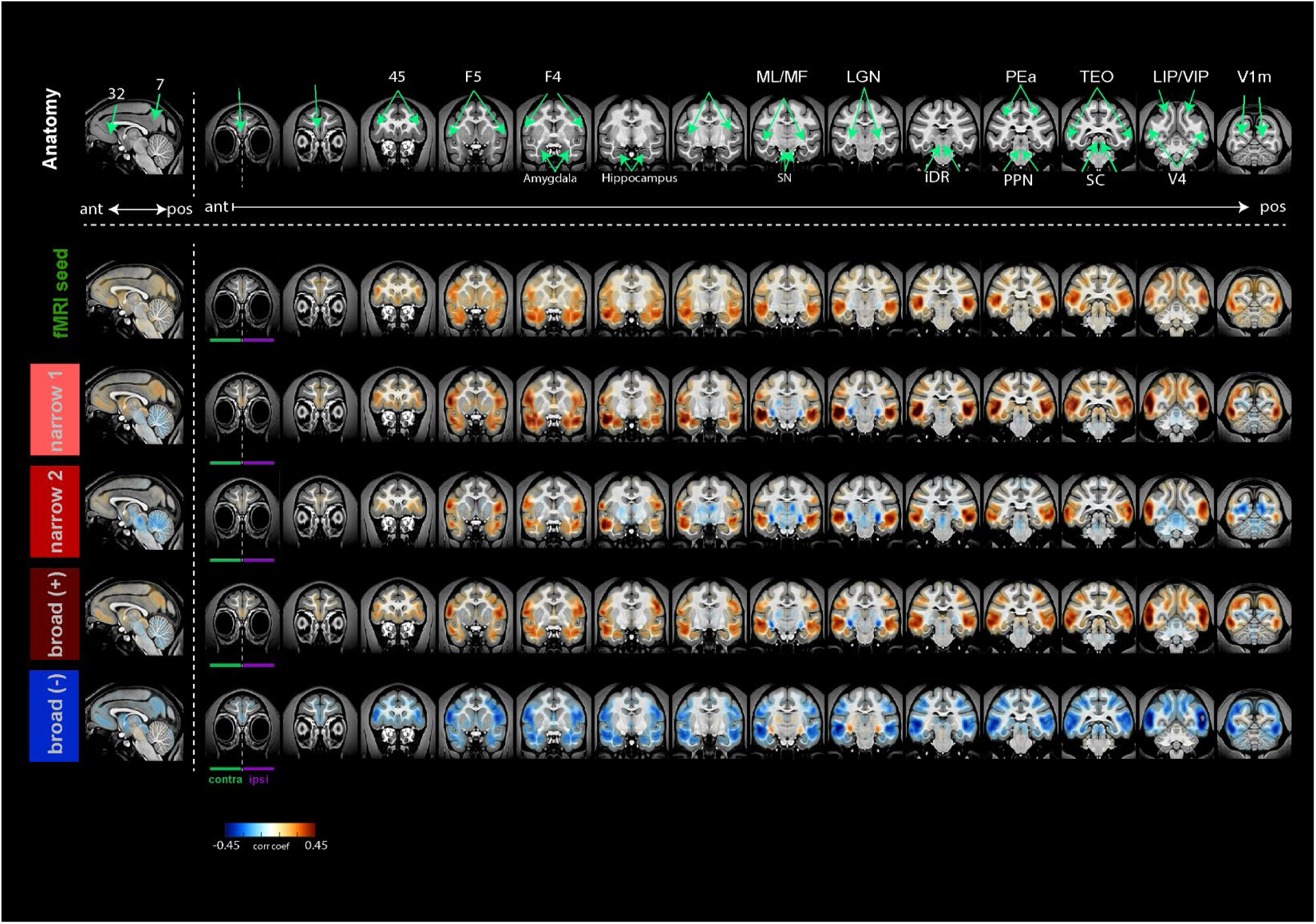
fMRI activation across groups of cells. Functional connetivity across the different groups of neurons. The top panel illustrates anatomical slices, with green arrows marking the anatomical locations of specific brain regions. The bottom panel depicts the functional connectivity across neuronal groups.

**Figure S8.**
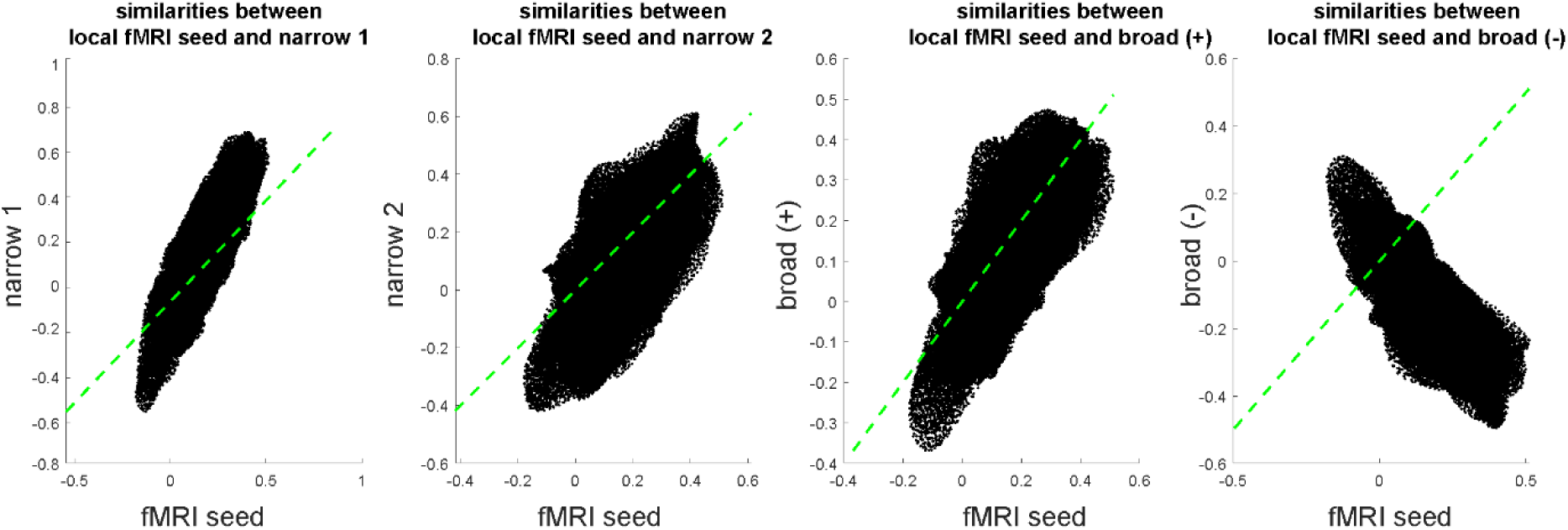
Voxel-wise similarity between the local fMRI seed map and each fMRI-cell group across the entire brain. Each panel shows a scatter plot comparing voxel-wise values from the fMRI seed to the corresponding voxel values from one cell-group map (narrow 1, narrow 2, broad (+), broad (–). Points represent all brain voxels, including both cortical and subcortical regions. Solid colored lines indicate the best-fit linear regression for each comparison, and the green dashed line denotes the identity line. Differences in slope and spread reflect how strongly each cell-group map aligns with the spatial pattern of the fMRI seed at the voxel level.

**Figure S9.**
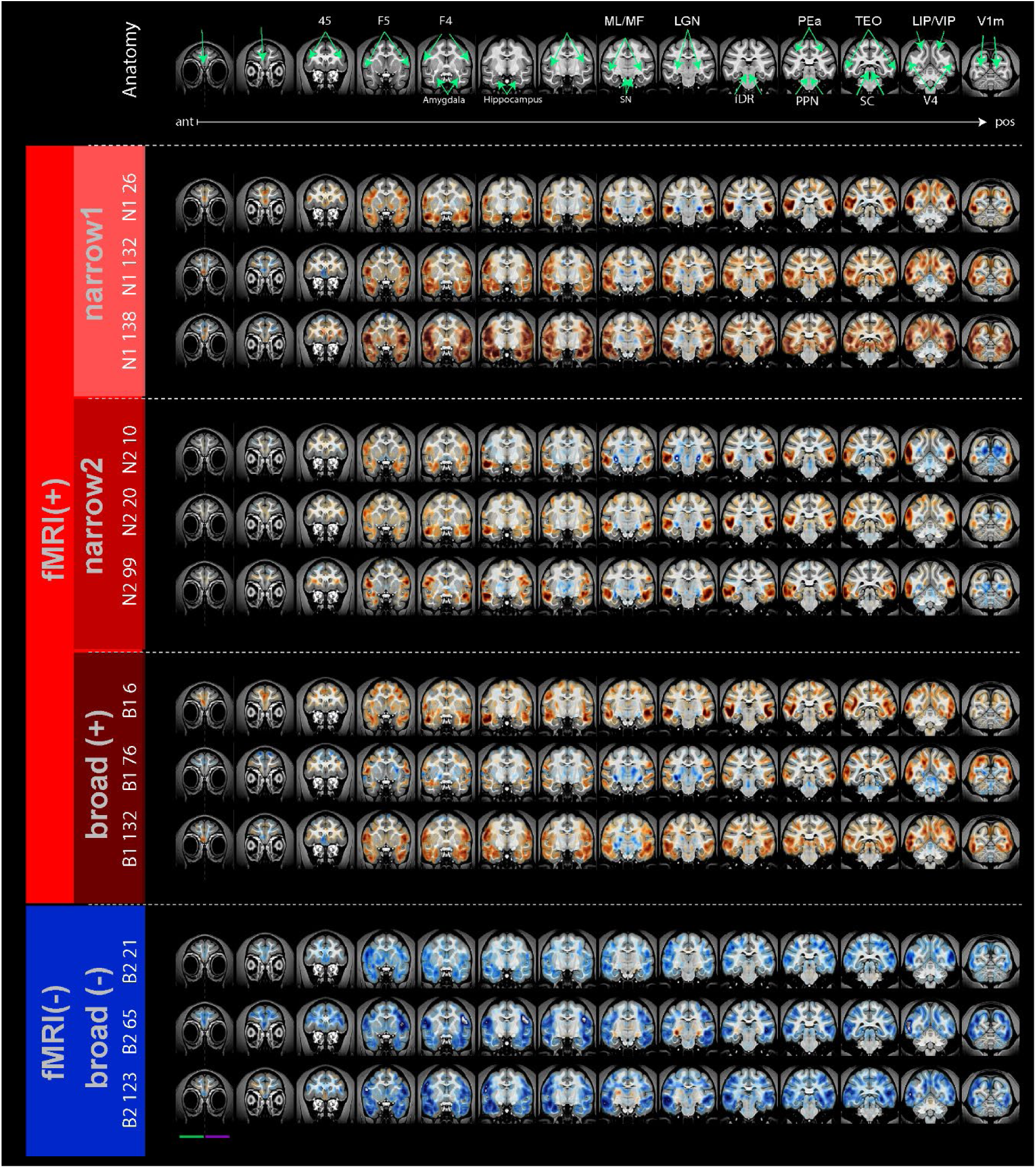
Example of single-unit fMRI maps. Each panel shows three representative fMRI maps derived from individual neurons, grouped according to their waveform-based classification (narrow 1, narrow 2, broad (+), broad (-). These examples illustrate the range and spatial patterns of whole-brain functional connectivity associated with single-unit activity within each neuronal subgroup.

**Figure S10.**
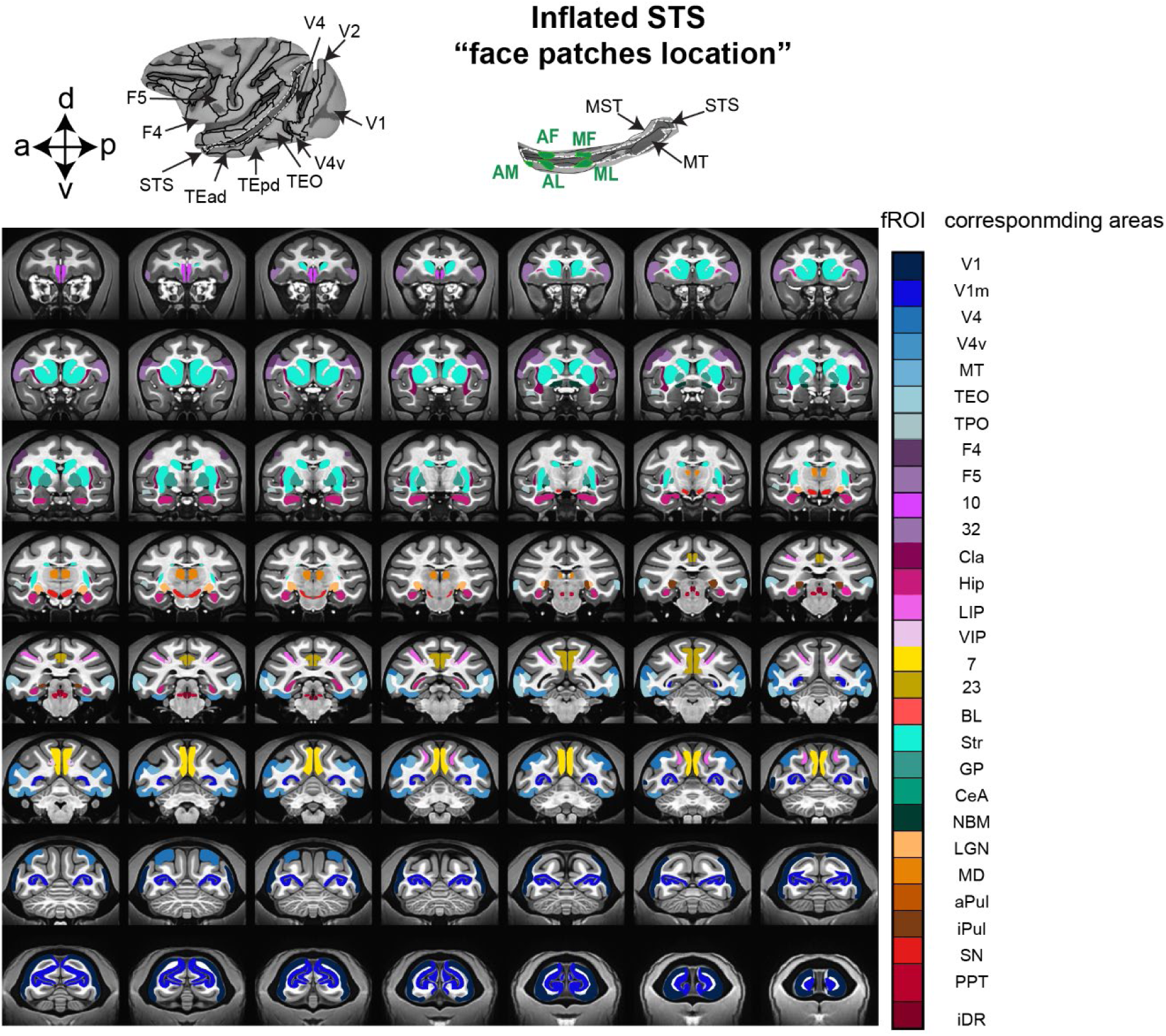
Functional regions of interest. Boundaries of the fROIS that were utilized in our current study.

## References

Acero-Pousa, I., Escrichs, A., Clara Dagnino, P., Sanz Perl, Y., Kringelbach, M.L., Uhlhaas, P.J., and Deco, G. (2025). Reconfiguration of functional brain hierarchy in schizophrenia. Transl Psychiatry 15, 356. 10.1038/s41398-025-03584-0.

Anenberg, E., Chan, A.W., Xie, Y., LeDue, J.M., and Murphy, T.H. (2015). Optogenetic stimulation of GABA neurons can decrease local neuronal activity while increasing cortical blood flow. J Cereb Blood Flow Metab 35, 1579–1586. 10.1038/jcbfm.2015.140.

Arsenault, J.T., Caspari, N., Vandenberghe, R., and Vanduffel, W. (2018). Attention Shifts Recruit the Monkey Default Mode Network. J Neurosci 38, 1202–1217. 10.1523/JNEUROSCI.1111-17.2017.

Bajada, C.J., Costa Campos, L.Q., Caspers, S., Muscat, R., Parker, G.J.M., Lambon Ralph, M.A., Cloutman, L.L., and Trujillo-Barreto, N.J. (2020). A tutorial and tool for exploring feature similarity gradients with MRI data. Neuroimage 221, 117140. 10.1016/j.neuroimage.2020.117140.

Bartho, P., Hirase, H., Monconduit, L., Zugaro, M., Harris, K.D., and Buzsaki, G. (2004). Characterization of neocortical principal cells and interneurons by network interactions and extracellular features. J Neurophysiol 92, 600–608. 10.1152/jn.01170.2003.

Barzo, P., Szots, I., Toth, M., Csajbok, E.A., Molnar, G., and Tamas, G. (2025). Electrophysiology and morphology of human cortical supragranular pyramidal cells in a wide age range. Elife 13, RP100390. 10.7554/eLife.100390.

Biswal, B., Yetkin, F.Z., Haughton, V.M., and Hyde, J.S. (1995a). Functional connectivity in the motor cortex of resting human brain using echo-planar MRI. Magn Reson Med 34, 537–541. 10.1002/mrm.1910340409.

Biswal, B., Yetkin, F.Z., Haughton, V.M., and Hyde, J.S. (1995b). Functional connectivity in the motor cortex of resting human brain using echo-planar MRI. Magnetic resonance in medicine : official journal of the Society of Magnetic Resonance in Medicine / Society of Magnetic Resonance in Medicine 34, 537–541.

Biswal, B.B., Mennes, M., Zuo, X.-N., Gohel, S., Kelly, C., Smith, S.M., Beckmann, C.F., Adelstein, J.S., Buckner, R.L., Colcombe, S., et al. (2010). Toward discovery science of human brain function. Proceedings of the National Academy of Sciences 107, 4734–4739. 10.1073/pnas.0911855107.

Bondar, I.V., Leopold, D.A., Richmond, B.J., Victor, J.D., and Logothetis, N.K. (2009). Long-term stability of visual pattern selective responses of monkey temporal lobe neurons. Plos One 4, e8222. 10.1371/journal.pone.0008222.

Buxton, R.B., Uludag, K., Dubowitz, D.J., and Liu, T.T. (2004). Modeling the hemodynamic response to brain activation. Neuroimage 23 *Suppl 1*, S220–233. 10.1016/j.neuroimage.2004.07.013.

Cardin, J.A., Carlen, M., Meletis, K., Knoblich, U., Zhang, F., Deisseroth, K., Tsai, L.H., and Moore, C.I. (2010). Targeted optogenetic stimulation and recording of neurons in vivo using cell-type-specific expression of Channelrhodopsin-2. Nat Protoc 5, 247–254. 10.1038/nprot.2009.228.

Coletta, L., Pagani, M., Whitesell, J.D., Harris, J.A., Bernhardt, B., and Gozzi, A. (2020). Network structure of the mouse brain connectome with voxel resolution. Sci Adv 6. 10.1126/sciadv.abb7187.

Connors, B.W., and Gutnick, M.J. (1990). Intrinsic firing patterns of diverse neocortical neurons. Trends Neurosci 13, 99–104. 10.1016/0166-2236(90)90185-d.

Cox, R.W. (1996). AFNI: software for analysis and visualization of functional magnetic resonance neuroimages. Comput Biomed Res 29, 162–173. 10.1006/cbmr.1996.0014.

Cruttenden, C.E., Zhu, W., Zhang, Y., Zhu, X.H., Chen, W., and Rajamani, R. (2022). Toward Completely Sampled Extracellular Neural Recording During fMRI. IEEE Trans Med Imaging 41, 1735–1746. 10.1109/TMI.2022.3149002.

Cutts, S.A., Faskowitz, J., Betzel, R.F., and Sporns, O. (2023). Uncovering individual differences in fine-scale dynamics of functional connectivity. Cereb Cortex 33, 2375–2394. 10.1093/cercor/bhac214.

Dearnley, B., Jones, M., Dervinis, M., and Okun, M. (2023). Brain state transitions primarily impact the spontaneous rate of slow-firing neurons. Cell Rep 42, 113185. 10.1016/j.celrep.2023.113185.

Deco, G., Jirsa, V.K., and McIntosh, A.R. (2011). Emerging concepts for the dynamical organization of resting-state activity in the brain. Nat Rev Neurosci 12, 43–56. 10.1038/nrn2961.

Ding, X., Froudist-Walsh, S., Jaramillo, J., Jiang, J., and Wang, X.J. (2024). Cell type-specific connectome predicts distributed working memory activity in the mouse brain. Elife 13. 10.7554/eLife.85442.

Du, Y., Fu, Z., and Calhoun, V.D. (2018). Classification and Prediction of Brain Disorders Using Functional Connectivity: Promising but Challenging. Front Neurosci 12, 525. 10.3389/fnins.2018.00525.

Elorette, C., Fujimoto, A., Stoll, F.M., Fujimoto, S.H., Bienkowska, N., London, L., Fleysher, L., Russ, B.E., and Rudebeck, P.H. (2024). The neural basis of resting-state fMRI functional connectivity in fronto-limbic circuits revealed by chemogenetic manipulation. Nat Commun 15, 4669. 10.1038/s41467-024-49140-0.

Fox, M.D., and Raichle, M.E. (2007). Spontaneous fluctuations in brain activity observed with functional magnetic resonance imaging. Nat Rev Neurosci 8, 700–711. 10.1038/nrn2201.

Friston, K.J. (2011). Functional and effective connectivity: a review. Brain Connectivity, 1–56. 10.1089/brain.2011.0008.

Fukushima, M., Saunders, R.C., Leopold, D.A., Mishkin, M., and Averbeck, B.B. (2012). Spontaneous high-gamma band activity reflects functional organization of auditory cortex in the awake macaque. Neuron 74, 899–910. 10.1016/j.neuron.2012.04.014.

Furlanis, E., Dai, M., Leyva Garcia, B., Tran, T., Vergara, J., Pereira, A., Gorissen, B.L., Wills, S., Vlachos, A., Hairston, A., et al. (2025). An enhancer-AAV toolbox to target and manipulate distinct interneuron subtypes. Neuron 113, 1525–1547.e1515. 10.1016/j.neuron.2025.05.002.

Goulas, A., Majka, P., Rosa, M.G.P., and Hilgetag, C.C. (2019). A blueprint of mammalian cortical connectomes. PLoS Biol 17, e2005346. 10.1371/journal.pbio.2005346.

Grimaldi, P., Saleem, K.S., and Tsao, D. (2016). Anatomical Connections of the Functionally Defined “Face Patches” in the Macaque Monkey. Neuron 90, 1325–1342. 10.1016/j.neuron.2016.05.009.

Gutierrez-Barragan, D., Ramirez, J.S.B., Panzeri, S., Xu, T., and Gozzi, A. (2024). Evolutionarily conserved fMRI network dynamics in the mouse, macaque, and human brain. Nat Commun 15, 8518. 10.1038/s41467-024-52721-8.

Gutierrez-Barragan, D., Singh, N.A., Alvino, F.G., Coletta, L., Rocchi, F., De Guzman, E., Galbusera, A., Uboldi, M., Panzeri, S., and Gozzi, A. (2022). Unique spatiotemporal fMRI dynamics in the awake mouse brain. Curr Biol 32, 631–644 e636. 10.1016/j.cub.2021.12.015.

He, B.J., Snyder, A.Z., Zempel, J.M., Smyth, M.D., and Raichle, M.E. (2008). Electrophysiological correlates of the brain’s intrinsic large-scale functional architecture. Proceedings of the National Academy of Sciences 105, 16039–16044. 10.1073/pnas.0807010105.

Henze, D.A., Borhegyi, Z., Csicsvari, J., Mamiya, A., Harris, K.D., and Buzsaki, G. (2000). Intracellular features predicted by extracellular recordings in the hippocampus in vivo. J Neurophysiol 84, 390–400. 10.1152/jn.2000.84.1.390.

Honey, C.J., Sporns, O., Cammoun, L., Gigandet, X., Thiran, J.P., Meuli, R., and Hagmann, P. (2009). Predicting human resting-state functional connectivity from structural connectivity. Proc Natl Acad Sci U S A 106, 2035–2040. 10.1073/pnas.0811168106.

Hong, S.J., Vos de Wael, R., Bethlehem, R.A.I., Lariviere, S., Paquola, C., Valk, S.L., Milham, M.P., Di Martino, A., Margulies, D.S., Smallwood, J., and Bernhardt, B.C. (2019). Atypical functional connectome hierarchy in autism. Nat Commun 10, 1022. 10.1038/s41467-019-08944-1.

Huang, Z.J., and Paul, A. (2019). The diversity of GABAergic neurons and neural communication elements. Nat Rev Neurosci 20, 563–572. 10.1038/s41583-019-0195-4.

Huber, L., Finn, E.S., Chai, Y., Goebel, R., Stirnberg, R., Stocker, T., Marrett, S., Uludag, K., Kim, S.G., Han, S., et al. (2021). Layer-dependent functional connectivity methods. Prog Neurobiol 207, 101835. 10.1016/j.pneurobio.2020.101835.

Hutchison, R.M., Leung, L.S., Mirsattari, S.M., Gati, J.S., Menon, R.S., and Everling, S. (2011). Resting-state networks in the macaque at 7T. NeuroImage 56, 1546–1555. 10.1016/j.neuroimage.2011.02.063.

Hutchison, R.M., Womelsdorf, T., Allen, E.A., Bandettini, P.A., Calhoun, V.D., Corbetta, M., Della Penna, S., Duyn, J.H., Glover, G.H., Gonzalez-Castillo, J., et al. (2013a). Dynamic functional connectivity: promise, issues, and interpretations. Neuroimage 80, 360–378. 10.1016/j.neuroimage.2013.05.079.

Hutchison, R.M., Womelsdorf, T., Allen, E.A., Bandettini, P.A., Calhoun, V.D., Corbetta, M., Della Penna, S., Duyn, J.H., Glover, G.H., Gonzalez-Castillo, J., et al. (2013b). Dynamic functional connectivity: promise, issues, and interpretations. NeuroImage 80, 360–378. 10.1016/j.neuroimage.2013.05.079.

Iraji, A., Fu, Z., Faghiri, A., Duda, M., Chen, J., Rachakonda, S., DeRamus, T., Kochunov, P., Adhikari, B.M., Belger, A., et al. (2023). Identifying canonical and replicable multi-scale intrinsic connectivity networks in 100k+ resting-state fMRI datasets. Hum Brain Mapp 44, 5729–5748. 10.1002/hbm.26472.

Isaacson, J.S., and Scanziani, M. (2011). How inhibition shapes cortical activity. Neuron 72, 231–243. 10.1016/j.neuron.2011.09.027.

Jang, H.J., Chung, H., Rowland, J.M., Richards, B.A., Kohl, M.M., and Kwag, J. (2020). Distinct roles of parvalbumin and somatostatin interneurons in gating the synchronization of spike times in the neocortex. Sci Adv 6, eaay5333. 10.1126/sciadv.aay5333.

Jung, Y.J., Sun, S.H., Almasi, A., Yunzab, M., Meffin, H., and Ibbotson, M.R. (2023). Characterization of extracellular spike waveforms recorded in wallaby primary visual cortex. Front Neurosci 17, 1244952. 10.3389/fnins.2023.1244952.

Kapogiannis, D., Reiter, D.A., Willette, A.A., and Mattson, M.P. (2013). Posteromedial cortex glutamate and GABA predict intrinsic functional connectivity of the default mode network. NeuroImage 64, 112–119. 10.1016/j.neuroimage.2012.09.029.

Kaufman, M.T., Churchland, M.M., and Shenoy, K.V. (2013). The roles of monkey M1 neuron classes in movement preparation and execution. J Neurophysiol 110, 817–825. 10.1152/jn.00892.2011.

Khader, P., Schicke, T., Roder, B., and Rosler, F. (2008). On the relationship between slow cortical potentials and BOLD signal changes in humans. Int J Psychophysiol 67, 252–261. 10.1016/j.ijpsycho.2007.05.018.

Kodama, N.X., Feng, T., Ullett, J.J., Chiel, H.J., Sivakumar, S.S., and Galan, R.F. (2018). Anti-correlated cortical networks arise from spontaneous neuronal dynamics at slow timescales. Sci Rep 8, 666. 10.1038/s41598-017-18097-0.

Koyano, K.W., Esch, E.M., Hong, J.J., Waidmann, E.N., Wu, H., and Leopold, D.A. (2023). Progressive neuronal plasticity in primate visual cortex during stimulus familiarization. Sci Adv 9, eade4648. 10.1126/sciadv.ade4648.

Koyano, K.W., Jones, A.P., McMahon, D.B.T., Waidmann, E.N., Russ, B.E., and Leopold, D.A. (2021). Dynamic Suppression of Average Facial Structure Shapes Neural Tuning in Three Macaque Face Patches. Curr Biol 31, 1–12 e15. 10.1016/j.cub.2020.09.070.

Krawchuk, M.B., Ruff, C.F., Yang, X., Ross, S.E., and Vazquez, A.L. (2020). Optogenetic assessment of VIP, PV, SOM and NOS inhibitory neuron activity and cerebral blood flow regulation in mouse somato-sensory cortex. Journal of Cerebral Blood Flow & Metabolism *40*, 1427–1440. 10.1177/0271678x19870105.

Landemard, A., Krumin, M., Harris, K.D., and Carandini, M. (2026). Brainwide blood volume reflects opposing neural populations. Nature. 10.1038/s41586-026-10350-9.

Lawrence, S.J.D., Formisano, E., Muckli, L., and de Lange, F.P. (2019). Laminar fMRI: Applications for cognitive neuroscience. Neuroimage 197, 785–791. 10.1016/j.neuroimage.2017.07.004.

Lee, E.K., Balasubramanian, H., Tsolias, A., Anakwe, S.U., Medalla, M., Shenoy, K.V., and Chandrasekaran, C. (2021). Non-linear dimensionality reduction on extracellular waveforms reveals cell type diversity in premotor cortex. Elife 10. 10.7554/eLife.67490.

Lee, J.H., Durand, R., Gradinaru, V., Zhang, F., Goshen, I., Kim, D.S., Fenno, L.E., Ramakrishnan, C., and Deisseroth, K. (2010). Global and local fMRI signals driven by neurons defined optogenetically by type and wiring. Nature 465, 788–792. 10.1038/nature09108.

Lee, S.H., Kwan, A.C., Zhang, S., Phoumthipphavong, V., Flannery, J.G., Masmanidis, S.C., Taniguchi, H., Huang, Z.J., Zhang, F., Boyden, E.S., et al. (2012). Activation of specific interneurons improves V1 feature selectivity and visual perception. Nature 488, 379–383. 10.1038/nature11312.

Leite, F.P., Tsao, D., Vanduffel, W., Fize, D., Sasaki, Y., Wald, L.L., Dale, A.M., Kwong, K.K., Orban, G.A., Rosen, B.R., et al. (2002). Repeated fMRI using iron oxide contrast agent in awake, behaving macaques at 3 Tesla. Neuroimage 16, 283–294. 10.1006/nimg.2002.1110.

Leopold, D.A., and Maier, A. (2011). Ongoing physiological processes in the cerebral cortex. NeuroImage. 10.1016/j.neuroimage.2011.10.059.

Leopold, D.A., Murayama, Y., and Logothetis, N.K. (2003). Very slow activity fluctuations in monkey visual cortex: implications for functional brain imaging. Cereb Cortex 13, 422–433. 10.1093/cercor/13.4.422.

Li, J.M., Acland, B.T., Brenner, A.S., Bentley, W.J., and Snyder, L.H. (2022). Relationships between correlated spikes, oxygen and LFP in the resting-state primate. Neuroimage 247, 118728. 10.1016/j.neuroimage.2021.118728.

Liu, C., Yen, C.C., Szczupak, D., Ye, F.Q., Leopold, D.A., and Silva, A.C. (2019). Anatomical and functional investigation of the marmoset default mode network. Nat Commun 10, 1975. 10.1038/s41467-019-09813-7.

Liu, X., de Zwart, J.A., Scholvinck, M.L., Chang, C., Ye, F.Q., Leopold, D.A., and Duyn, J.H. (2018). Subcortical evidence for a contribution of arousal to fMRI studies of brain activity. Nat Commun 9, 395. 10.1038/s41467-017-02815-3.

Liu, X., Leopold, D.A., and Yang, Y. (2021). Single-neuron firing cascades underlie global spontaneous brain events. Proc Natl Acad Sci U S A 118. 10.1073/pnas.2105395118.

Logothetis, N.K. (2008). What we can do and what we cannot do with fMRI. Nature 453, 869–878. 10.1038/nature06976.

Logothetis, N.K. (2015). Neural-Event-Triggered fMRI of large-scale neural networks. Curr Opin Neurobiol 31, 214–222. 10.1016/j.conb.2014.11.009.

Logothetis, N.K., Pauls, J., Augath, M., Trinath, T., and Oeltermann, A. (2001). Neurophysiological investigation of the basis of the fMRI signal. Nature 412, 150–157. 10.1038/35084005.

Lozano-Montes, L., Dimanico, M., Mazloum, R., Li, W., Nair, J., Kintscher, M., Schneggenburger, R., Harvey, M., and Rainer, G. (2020). Optogenetic Stimulation of Basal Forebrain Parvalbumin Neurons Activates the Default Mode Network and Associated Behaviors. Cell Rep 33, 108359. 10.1016/j.celrep.2020.108359.

Ma, X., Miraucourt, L.S., Qiu, H., Xu, M., Cook, E.P., Krishnaswamy, A., Sharif-Naeini, R., and Khadra, A. (2024). ElecFeX is a user-friendly toolbox for efficient feature extraction from single-cell electrophysiological recordings. Cell Rep Methods 4, 100791. 10.1016/j.crmeth.2024.100791.

Mantini, D., Gerits, A., Nelissen, K., Durand, J.B., Joly, O., Simone, L., Sawamura, H., Wardak, C., Orban, G.A., Buckner, R.L., and Vanduffel, W. (2011). Default mode of brain function in monkeys. J Neurosci 31, 12954–12962. 10.1523/JNEUROSCI.2318-11.2011.

Marik, S.A., Yamahachi, H., McManus, J.N., Szabo, G., and Gilbert, C.D. (2010). Axonal dynamics of excitatory and inhibitory neurons in somatosensory cortex. PLoS Biol 8, e1000395. 10.1371/journal.pbio.1000395.

Markicevic, M., Fulcher, B.D., Lewis, C., Helmchen, F., Rudin, M., Zerbi, V., and Wenderoth, N. (2020). Cortical Excitation:Inhibition Imbalance Causes Abnormal Brain Network Dynamics as Observed in Neurodevelopmental Disorders. Cerebral Cortex 30, 4922–4937. 10.1093/cercor/bhaa084.

Matsui, T., Murakami, T., and Ohki, K. (2019). Neuronal Origin of the Temporal Dynamics of Spontaneous BOLD Activity Correlation. Cereb Cortex 29, 1496–1508. 10.1093/cercor/bhy045.

Matsui, T., Tamura, K., Koyano, K.W., Takeuchi, D., Adachi, Y., Osada, T., and Miyashita, Y. (2011). Direct comparison of spontaneous functional connectivity and effective connectivity measured by intracortical microstimulation: an fMRI study in macaque monkeys. Cereb Cortex 21, 2348–2356. 10.1093/cercor/bhr019.

Milicevic, K.D., Barbeau, B.L., Lovic, D.D., Patel, A.A., Ivanova, V.O., and Antic, S.D. (2024). Physiological features of parvalbumin-expressing GABAergic interneurons contributing to high-frequency oscillations in the cerebral cortex. Current Research in Neurobiology 6, 100121. 10.1016/j.crneur.2023.100121.

Moeller, S., Freiwald, W.A., and Tsao, D.Y. (2008). Patches with links: a unified system for processing faces in the macaque temporal lobe. Science 320, 1355–1359. 10.1126/science.1157436.

Moon, H.S., Jiang, H., Vo, T.T., Jung, W.B., Vazquez, A.L., and Kim, S.G. (2021). Contribution of Excitatory and Inhibitory Neuronal Activity to BOLD fMRI. Cereb Cortex 31, 4053–4067. 10.1093/cercor/bhab068.

Moon, H.S., Vo, T.T., Ho Im, G., Hong, S.-J., and Kim, S.-G. (2025). Interhemispheric resting-state functional connectivity correlates with spontaneous neural interactions. Proceedings of the National Academy of Sciences 122, e2505294122. doi:10.1073/pnas.2505294122.

Murphy, K., Birn, R.M., Handwerker, D.A., Jones, T.B., and Bandettini, P.A. (2009). The impact of global signal regression on resting state correlations: are anti-correlated networks introduced? Neuroimage 44, 893–905. 10.1016/j.neuroimage.2008.09.036.

Nair, J., Klaassen, A.L., Arato, J., Vyssotski, A.L., Harvey, M., and Rainer, G. (2018). Basal forebrain contributes to default mode network regulation. Proc Natl Acad Sci U S A 115, 1352–1357. 10.1073/pnas.1712431115.

Naskar, S., Qi, J., Pereira, F., Gerfen, C.R., and Lee, S. (2021). Cell-type-specific recruitment of GABAergic interneurons in the primary somatosensory cortex by long-range inputs. Cell Rep 34, 108774. 10.1016/j.celrep.2021.108774.

Niazy, R.K., Beckmann, C.F., Iannetti, G.D., Brady, J.M., and Smith, S.M. (2005). Removal of FMRI environment artifacts from EEG data using optimal basis sets. Neuroimage 28, 720–737. 10.1016/j.neuroimage.2005.06.067.

Onorato, I., Tzanou, A., Schneider, M., Uran, C., Broggini, A.C., and Vinck, M. (2025). Distinct roles of PV and Sst interneurons in visually induced gamma oscillations. Cell Rep 44, 115385. 10.1016/j.celrep.2025.115385.

Pagani, M., Gutierrez-Barragan, D., de Guzman, A.E., Xu, T., and Gozzi, A. (2023). Mapping and comparing fMRI connectivity networks across species. Communications Biology 6, 1238. 10.1038/s42003-023-05629-w.

Petersen, P.C., Siegle, J.H., Steinmetz, N.A., Mahallati, S., and Buzsaki, G. (2021). CellExplorer: A framework for visualizing and characterizing single neurons. Neuron 109, 3594–3608 e3592. 10.1016/j.neuron.2021.09.002.

Peyrache, A., Dehghani, N., Eskandar, E.N., Madsen, J.R., Anderson, W.S., Donoghue, J.A., Hochberg, L.R., Halgren, E., Cash, S.S., and Destexhe, A. (2012). Spatiotemporal dynamics of neocortical excitation and inhibition during human sleep. Proc Natl Acad Sci U S A 109, 1731–1736. 10.1073/pnas.1109895109.

Pijnenburg, R., Scholtens, L.H., Mantini, D., Vanduffel, W., Barrett, L.F., and van den Heuvel, M.P. (2019). Biological Characteristics of Connection-Wise Resting-State Functional Connectivity Strength. Cerebral Cortex 29, 4646–4653. 10.1093/cercor/bhy342.

Quiroga, R.Q., Nadasdy, Z., and Ben-Shaul, Y. (2004). Unsupervised spike detection and sorting with wavelets and superparamagnetic clustering. Neural Comput 16, 1661–1687. 10.1162/089976604774201631.

Rocchi, F., Canella, C., Noei, S., Gutierrez-Barragan, D., Coletta, L., Galbusera, A., Stuefer, A., Vassanelli, S., Pasqualetti, M., Iurilli, G., et al. (2022). Increased fMRI connectivity upon chemogenetic inhibition of the mouse prefrontal cortex. Nature Communications 13, 1056. 10.1038/s41467-022-28591-3.

Russ, B.E., Koyano, K.W., Day-Cooney, J., Perwez, N., and Leopold, D.A. (2023). Temporal continuity shapes visual responses of macaque face patch neurons. Neuron 111, 903–914 e903. 10.1016/j.neuron.2022.12.021.

Saggar, M., Sporns, O., Gonzalez-Castillo, J., Bandettini, P.A., Carlsson, G., Glover, G., and Reiss, A.L. (2018). Towards a new approach to reveal dynamical organization of the brain using topological data analysis. Nat Commun 9, 1399. 10.1038/s41467-018-03664-4.

Sastre-Yagüe, D., Blanco Malerba, S., Rocchi, F., Gini, S., Mancini, G., Stuefer, A., Coletta, L., Noei, S., Markicevic, M., Alvino, F.G., et al. (2026). Cortical excitability inversely modulates fMRI connectivity via low-frequency neuronal coupling. bioRxiv, 2026.2003.2012.710517. 10.64898/2026.03.12.710517.

Schmidt, M., Bakker, R., Shen, K., Bezgin, G., Diesmann, M., and van Albada, S.J. (2018). A multi-scale layer-resolved spiking network model of resting-state dynamics in macaque visual cortical areas. PLOS Computational Biology 14, e1006359. 10.1371/journal.pcbi.1006359.

Schölvinck, M.L., Leopold, D.A., Brookes, M.J., and Khader, P.H. (2013). The contribution of electrophysiology to functional connectivity mapping. NeuroImage 80, 297–306. 10.1016/j.neuroimage.2013.04.010.

Schölvinck, M.L., Maier, A., Ye, F.Q., Duyn, J.H., and Leopold, D.A. (2010). Neural basis of global resting-state fMRI activity. Proceedings of the National Academy of Sciences 107, 10238–10243. 10.1073/pnas.0913110107.

Seidlitz, J., Sponheim, C., Glen, D., Ye, F.Q., Saleem, K.S., Leopold, D.A., Ungerleider, L., and Messinger, A. (2018). A population MRI brain template and analysis tools for the macaque. Neuroimage 170, 121–131. 10.1016/j.neuroimage.2017.04.063.

Shahsavarani, S., Thibodeaux, D.N., Xu, W., Kim, S.H., Lodgher, F., Nwokeabia, C., Cambareri, M., Yagielski, A.J., Zhao, H.T., Handwerker, D.A., et al. (2023). Cortex-wide neural dynamics predict behavioral states and provide a neural basis for resting-state dynamic functional connectivity. Cell Rep 42, 112527. 10.1016/j.celrep.2023.112527.

Smirnakis, S.M., Schmid, M.C., Weber, B., Tolias, A.S., Augath, M., and Logothetis, N.K. (2007). Spatial specificity of BOLD versus cerebral blood volume fMRI for mapping cortical organization. J Cereb Blood Flow Metab 27, 1248–1261. 10.1038/sj.jcbfm.9600434.

Smitha, K.A., Arun, K.M., Rajesh, P.G., Thomas, B., and Kesavadas, C. (2017). Resting-State Seed-Based Analysis: An Alternative to Task-Based Language fMRI and Its Laterality Index. AJNR Am J Neuroradiol 38, 1187–1192. 10.3174/ajnr.A5169.

Snyder, A.C., Morais, M.J., and Smith, M.A. (2016). Dynamics of excitatory and inhibitory networks are differentially altered by selective attention. J Neurophysiol 116, 1807–1820. 10.1152/jn.00343.2016.

Sporns, O., Faskowitz, J., Teixeira, A.S., Cutts, S.A., and Betzel, R.F. (2021). Dynamic expression of brain functional systems disclosed by fine-scale analysis of edge time series. Netw Neurosci 5, 405–433. 10.1162/netn_a_00182.

Stagg, C.J., Bachtiar, V., Amadi, U., Gudberg, C.A., Ilie, A.S., Sampaio-Baptista, C., O’Shea, J., Woolrich, M., Smith, S.M., Filippini, N., et al. (2014). Local GABA concentration is related to network-level resting functional connectivity. eLife 3, e01465. 10.7554/eLife.01465.

Stringer, C., Pachitariu, M., Steinmetz, N., Reddy, C.B., Carandini, M., and Harris, K.D. (2019). Spontaneous behaviors drive multidimensional, brainwide activity. Science 364, 255. 10.1126/science.aav7893.

Sundqvist, N., Podéus, H., Sten, S., Engström, M., Dura-Bernal, S., and Cedersund, G. (2025). Model-driven meta-analysis establishes a new consensus view: Inhibitory neurons dominate BOLD-fMRI responses. Computers in Biology and Medicine 197, 111014. 10.1016/j.compbiomed.2025.111014.

Taylor, J.J., Kurt, H.G., and Anand, A. (2021). Resting State Functional Connectivity Biomarkers of Treatment Response in Mood Disorders: A Review. Front Psychiatry 12, 565136. 10.3389/fpsyt.2021.565136.

Torres-Gomez, S., Blonde, J.D., Mendoza-Halliday, D., Kuebler, E., Everest, M., Wang, X.J., Inoue, W., Poulter, M.O., and Martinez-Trujillo, J. (2020). Changes in the Proportion of Inhibitory Interneuron Types from Sensory to Executive Areas of the Primate Neocortex: Implications for the Origins of Working Memory Representations. Cereb Cortex 30, 4544–4562. 10.1093/cercor/bhaa056.

Trainito, C., von Nicolai, C., Miller, E.K., and Siegel, M. (2019). Extracellular Spike Waveform Dissociates Four Functionally Distinct Cell Classes in Primate Cortex. Curr Biol 29, 2973–2982 e2975. 10.1016/j.cub.2019.07.051.

Tremblay, R., Lee, S., and Rudy, B. (2016). GABAergic Interneurons in the Neocortex: From Cellular Properties to Circuits. Neuron 91, 260–292. 10.1016/j.neuron.2016.06.033.

Tsodyks, M., Kenet, T., Grinvald, A., and Arieli, A. (1999). Linking spontaneous activity of single cortical neurons and the underlying functional architecture. Science 286, 1943–1946. 10.1126/science.286.5446.1943.

Turchi, J., Chang, C., Ye, F.Q., Russ, B.E., Yu, D.K., Cortes, C.R., Monosov, I.E., Duyn, J.H., and Leopold, D.A. (2018). The Basal Forebrain Regulates Global Resting-State fMRI Fluctuations. Neuron 97, 940–952.e944. 10.1016/j.neuron.2018.01.032.

Uhlirova, H., Kilic, K., Tian, P., Thunemann, M., Desjardins, M., Saisan, P.A., Sakadzic, S., Ness, T.V., Mateo, C., Cheng, Q., et al. (2016). Cell type specificity of neurovascular coupling in cerebral cortex. Elife 5. 10.7554/eLife.14315.

van den Brink, R.L., Pfeffer, T., Warren, C.M., Murphy, P.R., Tona, K.D., van der Wee, N.J., Giltay, E., van Noorden, M.S., Rombouts, S.A., Donner, T.H., and Nieuwenhuis, S. (2016). Catecholaminergic Neuromodulation Shapes Intrinsic MRI Functional Connectivity in the Human Brain. J Neurosci 36, 7865–7876. 10.1523/JNEUROSCI.0744-16.2016.

Vigneswaran, G., Kraskov, A., and Lemon, R.N. (2011). Large identified pyramidal cells in macaque motor and premotor cortex exhibit “thin spikes”: implications for cell type classification. J Neurosci 31, 14235–14242. 10.1523/JNEUROSCI.3142-11.2011.

Vincent, J.L., Patel, G.H., Fox, M.D., Snyder, A.Z., Baker, J.T., Van Essen, D.C., Zempel, J.M., Snyder, L.H., Corbetta, M., and Raichle, M.E. (2007). Intrinsic functional architecture in the anaesthetized monkey brain. Nature 447, 83–86. 10.1038/nature05758.

Vo, T.T., Im, G.H., Han, K., Suh, M., Drew, P.J., and Kim, S.G. (2023). Parvalbumin interneuron activity drives fast inhibition-induced vasoconstriction followed by slow substance P-mediated vasodilation. Proc Natl Acad Sci U S A 120, e2220777120. 10.1073/pnas.2220777120.

Vo, T.T., Jung, W.B., Jin, T., Im, G.H., Lee, S., and Kim, S.G. (2025). Somatostatin-expressing interneurons induce early NO-driven and late specific astrocyte-mediated vasodilation. Nat Commun 16, 6606. 10.1038/s41467-025-61771-5.

Wang, Z., Chen, L.M., Negyessy, L., Friedman, R.M., Mishra, A., Gore, J.C., and Roe, A.W. (2013). The relationship of anatomical and functional connectivity to resting-state connectivity in primate somatosensory cortex. Neuron 78, 1116–1126. 10.1016/j.neuron.2013.04.023.

Woodward, N.D., and Cascio, C.J. (2015). Resting-State Functional Connectivity in Psychiatric Disorders. JAMA Psychiatry 72, 743–744. 10.1001/jamapsychiatry.2015.0484.

Xiang, Q.S., and Ye, F.Q. (2007). Correction for geometric distortion and N/2 ghosting in EPI by phase labeling for additional coordinate encoding (PLACE). Magn Reson Med 57, 731–741. 10.1002/mrm.21187.

Yacoub, E., Grier, M.D., Auerbach, E.J., Lagore, R.L., Harel, N., Adriany, G., Zilverstand, A., Hayden, B.Y., Heilbronner, S.R., Ugurbil, K., and Zimmermann, J. (2020). Ultra-high field (10.5 T) resting state fMRI in the macaque. Neuroimage 223, 117349. 10.1016/j.neuroimage.2020.117349.

Yang, J., Gohel, S., and Vachha, B. (2020). Current methods and new directions in resting state fMRI. Clin Imaging 65, 47–53. 10.1016/j.clinimag.2020.04.004.

Young, N.A., Collins, C.E., and Kaas, J.H. (2013). Cell and neuron densities in the primary motor cortex of primates. Front Neural Circuits 7, 30. 10.3389/fncir.2013.00030.

Zaitsev, A.V., Povysheva, N.V., Gonzalez-Burgos, G., Rotaru, D., Fish, K.N., Krimer, L.S., and Lewis, D.A. (2009). Interneuron diversity in layers 2-3 of monkey prefrontal cortex. Cereb Cortex 19, 1597–1615. 10.1093/cercor/bhn198.

Zaldivar, D., Koyano, K.W., Ye, F.Q., Godlove, D.C., Park, S.H., Russ, B.E., Bhik-Ghanie, R., and Leopold, D.A. (2022). Brain-wide functional connectivity of face patch neurons during rest. Proc Natl Acad Sci U S A 119, e2206559119. 10.1073/pnas.2206559119.

Zaldivar, D., Rauch, A., Logothetis, N.K., and Goense, J. (2018). Two distinct profiles of fMRI and neurophysiological activity elicited by acetylcholine in visual cortex. Proc Natl Acad Sci U S A 115, E12073–E12082. 10.1073/pnas.1808507115.

